# Distinct trajectories in low-dimensional neural oscillation state space track dynamic decision-making in humans

**DOI:** 10.1101/2022.06.14.494674

**Authors:** Thomas Thiery, Pierre Rainville, Paul Cisek, Karim Jerbi

## Abstract

The brain evolved to govern behavior in a dynamic world, in which pertinent information about choices is often in flux. Thus, the commitment to an action choice must reflect a balance between monitoring that information and the necessity to act before opportunities are lost. Here, we investigate the mechanisms of dynamic decision-making in humans using low dimensional space representation of brain wide magnetoencephalography recordings. We show that the principal components (PCs) of alpha (9-13 Hz) and beta power (16-24 Hz) are involved in tracking sensory information evolving over time in the sensorimotor and visual cortex. We also found that alpha PCs reflect the commitment to a particular choice, while beta PCs reflect motor execution. Finally, higher frequency components in subcortical areas reflect the adjustment of speed-accuracy tradeoff policies. These results provide a new detailed characterization of the distributed oscillatory brain processes underlying dynamic decision-making in humans.

## INTRODUCTION

In our daily lives we make a wide variety of decisions. Some are abstract and long-term, such as choosing a career, and some are more mundane, such as whether to turn right or left at an intersection. In most cases there are risks, costs, and benefits to selecting a given course of action, and a rich body of work exists on how humans and other animals weigh these factors in making and committing to their choice. In particular, recent studies using functional MRI, electro-encephalography (EEG), and magnetoencephalography (MEG) have yielded important insights into the neural mechanisms of decision-making in humans^1–13^. For example, EEG and MEG studies have reported choice-predictive neuronal oscillations in motor, premotor and early visual cortex, thought to reflect accumulated sensory evidence^8–10,12–15^. While low frequency oscillations are thought to reflect global processing and feedforward information flow through the cortical hierarchy, high frequencies have been associated with local processing and feedback cortical pathways^16–23^. Beta oscillations (13-30 Hz) in particular are thought to play a crucial role in motor control and have been important for understanding the neural mechanisms underlying decisions between competing actions^24,25^.

While these studies are building a strong foundation for our understanding of the neural bases of decisions, many questions remain unanswered. Prominent among these is the question of how the brain makes the transition from deliberating about the pros and cons of given options to committing to, and executing, a single choice. One potential answer comes from the well known “drift-diffusion model” (DDM), which suggests that commitment occurs when sensory (or value-based) evidence for one choice over the others is accumulated to some threshold level that controls the speed-accuracy trade-off^26,27^, and at that point a response is chosen that is subsequently transformed into an action plan and executed. Consistent with this proposal, neural activity in many cortical and subcortical regions builds-up at a rate that is related to the evidence^28,29^. However, in most tasks studied to date, the evidence in favor of each given choice is constant over the course of a trial such that evidence accumulation reflects essentially the speed of perceptual processing/integration. An example is the widely used random dot motion discrimination task^30^, in which subjects make a decision about the direction of coherent motion in a field of randomly moving dots. However, in conditions where motion coherence is stable across a given trial, it is notoriously difficult to distinguish neural activity build-up that is due to evidence accumulation from build-up related to motor preparation, reward anticipation, the urge to make a choice, or simply the passage of time^31–36^.

One approach for distinguishing processes related to the deliberation from other processes like motor preparation is to use “dynamic decision-making tasks”. In such paradigms, sensory information in favor of competing responses varies over time within each trial. This makes it possible to isolate neural activity that follows the changing evidence from neural activity that is related to other processes, such as the preparation of a response after the choice is made. Such studies have suggested that the computation of sensory evidence is in fact quite fast, on the order of 200ms or so^37–39^, and that most of the build-up of neural activity may be caused by a rising “urgency signal” that pushes the system to commit to a choice even if the evidence is weak^31,32,40^. This mechanism, sometimes referred to as “collapsing bounds”, has been suggested as an effective policy for maximizing reward rates^41–45^. These effects question some of the assumptions of the DDM and call for a more refined account of distinctive processes contributing to decision making.

One variation, called the “urgency gating model” (UGM)^32,35,41^, suggests that evidence is computed quickly with a small time-constant of integration (emphasizing novel information), and combined with a context-dependent urgency signal that dynamically adjusts the threshold for initiating movement. While this model produces behavioral and neural predictions that are nearly indistinguishable from the DDM in tasks with constant evidence^32,35^, such as random dot motion discrimination, it makes very different predictions in tasks in which evidence is changing dynamically. In those conditions, results from numerous studies are better explained by the UGM than the DDM, at both a behavioral and neural level ^5,10,32,35,40,41,46–48,39^. Furthermore, a series of electrophysiological studies in monkeys suggests that fast estimates of sensory evidence are computed in the prefrontal cortex^49^, combined with an urgency signal from the basal ganglia^46^, and then used to bias a competition evolving within the dorsal premotor and primary motor cortex^40,49^. The urgency signal is adjusted based on context^34,50^, including prior trial history^51^, but always involves rising activity that drives the animal to eventually make a choice even if the evidence is ambiguous. In situations that require fast decisions, the urgency signal will be stronger and can be operationalized as shorter response time and increased error rates, consistent with classical speed-accuracy trade-off principles.

While these findings have illuminated the mechanistic basis of decisions in a variety of conditions in non-human primates, it is not clear to what extent they may be specific to highly over-trained monkeys^52^. Furthermore, local recordings do not inform about the large-scale distribution of these computations. In the present study, we address these outstanding issues by recording whole brain MEG activity of human subjects while they perform a dynamic decision-making task. We combine dimensionality reduction using Principal Component Analysis (PCA) and machine learning to identify spectral, spatial, and temporal features of neuronal activity specifically related to (i) tracking sensory evidence during deliberation, (ii) committing to a particular choice, (iii) motor preparation and execution and (iv) control over speed-accuracy.

## RESULTS

In the tokens task, participants (N=28) pressed a button with their right or left index fingers based on their guess about which of two peripheral targets would receive the majority of 15 tokens jumping from a central circle to one of the two targets every 200 ms (Figure 1A). Participants were free to decide at any time and asked to get as many correct guesses as possible, motivating them to optimize successes per unit time. After a target was chosen, the remaining tokens accelerated, jumping every 150 ms in “slow” blocks or every 50 ms in “fast” blocks (Figure 1B). The design of the task thus presented a speed versus accuracy trade-off (SAT): subjects could choose to wait until all tokens have jumped (i.e., conservative decisions), or to take a guess before all tokens have jumped (i.e., hasty decisions) and save time, with the risk of having made the wrong choice. Each subject completed eight blocks, alternating between fast blocks which encourage hasty decisions (more time is saved by guessing quickly), and slow blocks which encourage more conservative decisions. Although subjects were told that each token jump was completely random, we interspersed among them three specific categories of trials characterized on the basis of the temporal pattern of jumps: Easy, ambiguous, and misleading trials (see Figure 1C). At the end of each trial, feedback indicating correct or incorrect choices was presented to subjects (Figure 1A, red or green circles). Before and after the tokens task, subjects performed a two-choice visuo-motor reaction-time task (two 2-minute blocks), in which the 15 tokens originally arranged in the center circle all simultaneously jumped (“GO signal) into one of the two peripheral targets (left or right) after a variable delay. The mean reaction time obtained during the visuo-motor reaction-time task was taken as an estimate of processing delays not related to decision-making and used to estimate decision time for each trial of the tokens task (*Button Press Time* – *mean Reaction Time* = *Decision Time*, see Figure 1B). This experimental design allowed us to identify whole-brain dynamics involved in different aspects of decision-making, including the process of deliberation, the moment of commitment, and the adjustment of speed versus accuracy trade-offs.

**Figure 1.**
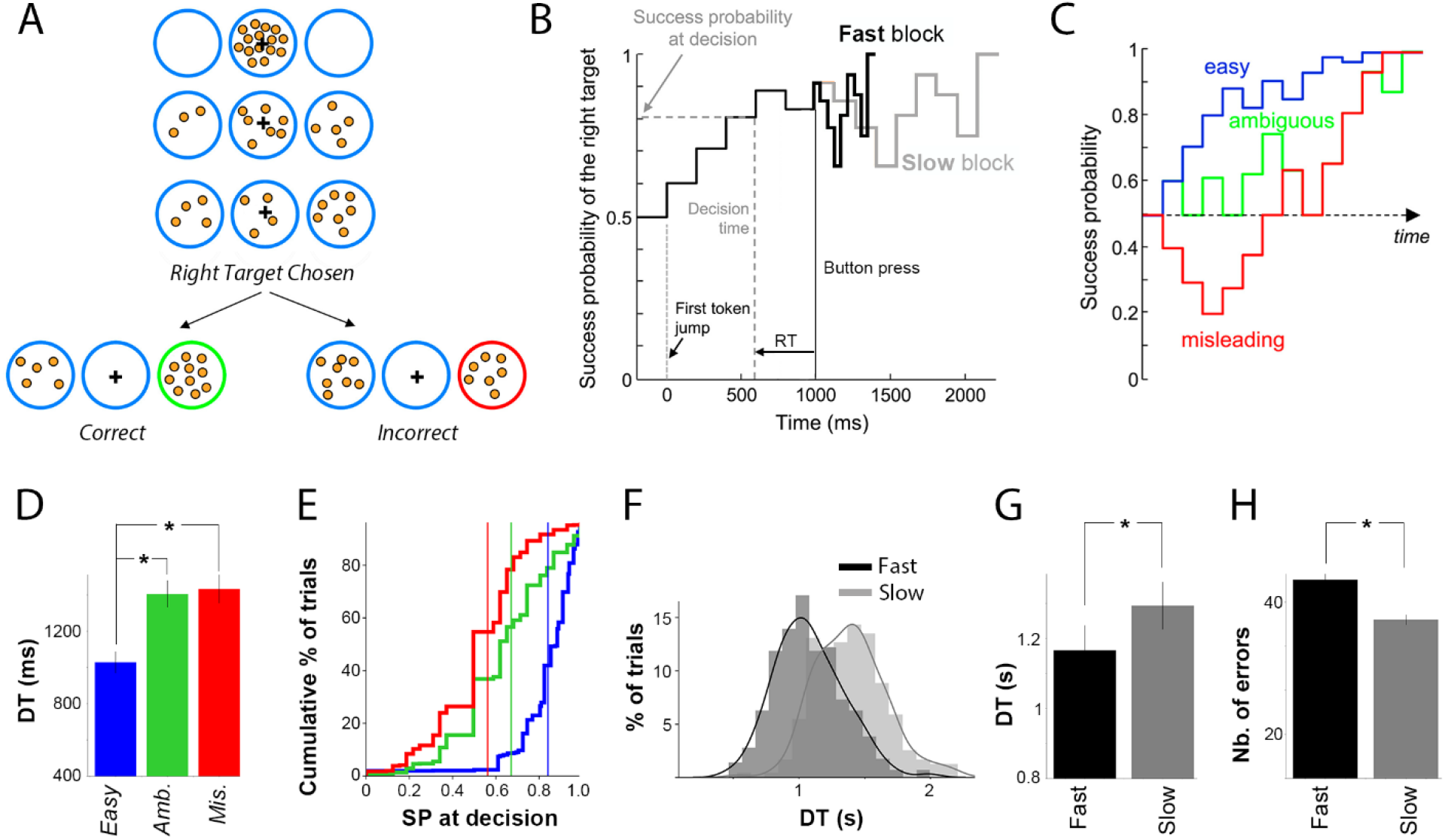
Task and behavior. **A**. The tokens task. Each row illustrates steps in an example trial, in which the right target is chosen. Two example outcomes are then illustrated, one in which the remaining tokens keep jumping to the right target (easy trial, correct choice), and one in which the remaining tokens jump to the left target (misleading trial, incorrect choice). **B**. Success probability (SP) profile of an example trial, in which the right target was chosen. The thin black trace indicates SP before the target is chosen, while tokens jump every 200 ms. After the target is chosen (button press, vertical solid line), the remaining tokens jump either every 150 ms (in slow blocks, grey trace) or every 50 ms (fast block, thick black trace). To yield an estimate of the decision time (DT, vertical gray dashed line), and the success probability at which the participant committed to his choice (horizontal gray dashed line), we subtract the mean RT calculated using a two-choice visuo-motor reaction-time task from button press time (horizontal black arrow). **C**. Example success probability profiles of “easy” (blue), “ambiguous” (green), or “misleading” (red) trials (see criteria in Methods). **D**. Mean decision time across subjects (N=28) in easy trials was significantly shorter than in ambiguous and misleading trials (* p<0.001). **E**. Cumulative distributions of success probabilities at decision time for easy (blue), ambiguous (green) and misleading (red) trials for all subjects. Vertical lines indicate mean success probability. **F**. Distributions of the decision times of a representative subject for fast (black), and slow (grey) blocks. **G**. The mean decision time across participants (N=28) in fast blocks was significantly shorter than in slow blocks. **H**. Participants made significantly more errors in fast blocks compared to slow blocks.

### Behavioural analysis

Decision time and success probability at the time of choice depended upon the sensory evidence provided by the distribution of tokens. Subjects made their decision significantly faster during easy trials (mean DT = 1028 ± 59 ms) than during both ambiguous (mean = 1405 ± 74 ms, Paired student t-test, easy *vs* ambiguous, *T value = -15*.*04; p = 1*.*19 e-14*) and misleading trials (mean = 1433 ± 79 ms, Paired student t-test, easy *vs* misleading, *T value = -13*.*1; p = 3*.*25 e-13*, Figure 1D), and their success probability at decision time was significantly higher in easy trials (Figure 1E). However, no significant differences were found between DT in ambiguous and misleading trials (Paired student t-test, ambiguous *vs* misleading, *T value = -1,84; p = 0*.*077)*. Subjects also behaved more hastily and made their decision earlier in the fast blocks (mean DT = 1166 ± 71 ms) compared to slow blocks (mean = 1293 ± 68 ms, Paired student t-test, *T value = -6,08; p = 1*.*71 e-6)*, during which they were more conservative. Moreover, participants made significantly more mistakes in fast blocks (mean = 45.1 ± 9.9 errors) compared to slow blocks (mean = 36 ± 6 errors; Paired student t-test, errors in fast *vs* slow blocks, *T value = 6*.*1; p = 1*.*7 e-6*), consistent with a trade-off between speed and accuracy. The distribution of DTs in fast and slow blocks for one subject, mean DTs and mean errors across subjects are shown in Figure 1F, G, H. Note that the present behavioral results replicate previous observations in both humans^32^ and rhesus monkeys^53^.

### State space analysis of neuronal oscillations

To study how modulations of neural activity dynamically reflect variables relevant to the performance of the tokens task we computed single-trial time-resolved spectral power envelopes in five frequency bands: theta (θ) [5–8 Hz], alpha (α) [9–13 Hz], beta (β) [16–24 Hz], gamma (γ) [30–60 Hz] and high gamma (high γ) [60–90 Hz] in the source space. The width of frequency bands was chosen based on results obtained with the FOOOF toolbox^54^, allowing to detect oscillations (peaks) in the frequency domain by separating the periodic and aperiodic components of neural power spectra (see Supplementary Figure S1 and Material and Methods). We then focussed our analyses of brain oscillations in a specific low-dimensional subspace that captures across-trial variance which arises from subjects’ choice (left or right choices), trial types (easy, ambiguous, or misleading trials), and block-dependent SAT policies (fast or slow blocks), similar to analyses of neural spiking activity in monkeys^49^. We computed this task-related subspace using principal component analysis (PCA, see Supplementary Figure S2 for an illustration). This yielded an unbiased estimate of the most prominent features (that is, patterns of neural activity) in whole brain sources across all subjects for each frequency band. We restricted subsequent analyses to the subspace spanned by the first 20 principal components (PCs), though our conclusions are mostly based on the top 4. Compared to more standard power analyses, the PCs generated can be thought of as the ‘building blocks’ of the observed neural activity, in that the activity of any specific source is a linear combination (weighted average) of these components across the twelve groups of trials (slow/fast blocks x left/right choices x easy/ambiguous/misleading) used for the PCA. For example, Supplementary Figure S3 shows the comparison between power envelopes and PCs generated for the same frequency band (beta) and the same ROI (sensorimotor cortex). Note that a conventional power analysis would find the well-known beta suppression and rebound, which is captured by PC1, but it would not reveal additional features such as choice dependence, captured by PC2. Furthermore, by applying the analysis simultaneously across all brain sites, we do not need to pre-define ROIs but can let the data reveal the regions that contribute most to these distinct components of oscillatory power, as will be shown below.

The first four PCs for theta, alpha, beta, and gamma oscillations explained most of the variance in activity over time (explaining 83.5%, 86.5%, 94%, and 76.7% respectively), and Figure 2 shows their temporal profiles after aligning the data to button presses (see Supplementary Figure S4 for the first 8 PCs and Supplementary Figure S5 for the same PCs aligned on the first token jump for each frequency band, including high gamma). Then, we used a Linear Discriminant Analysis (LDA) classifier to probe the ability of these spectral features to significantly decode the three task-related conditions of choice, trial class, and SAT policy in a data-driven manner. The statistical significance of time-resolved decoding accuracy was determined using permutation tests, corrected for multiple comparisons across subjects and time (*p < 0*.*05*, corrected across time, see Methods section).

**Figure 2.**
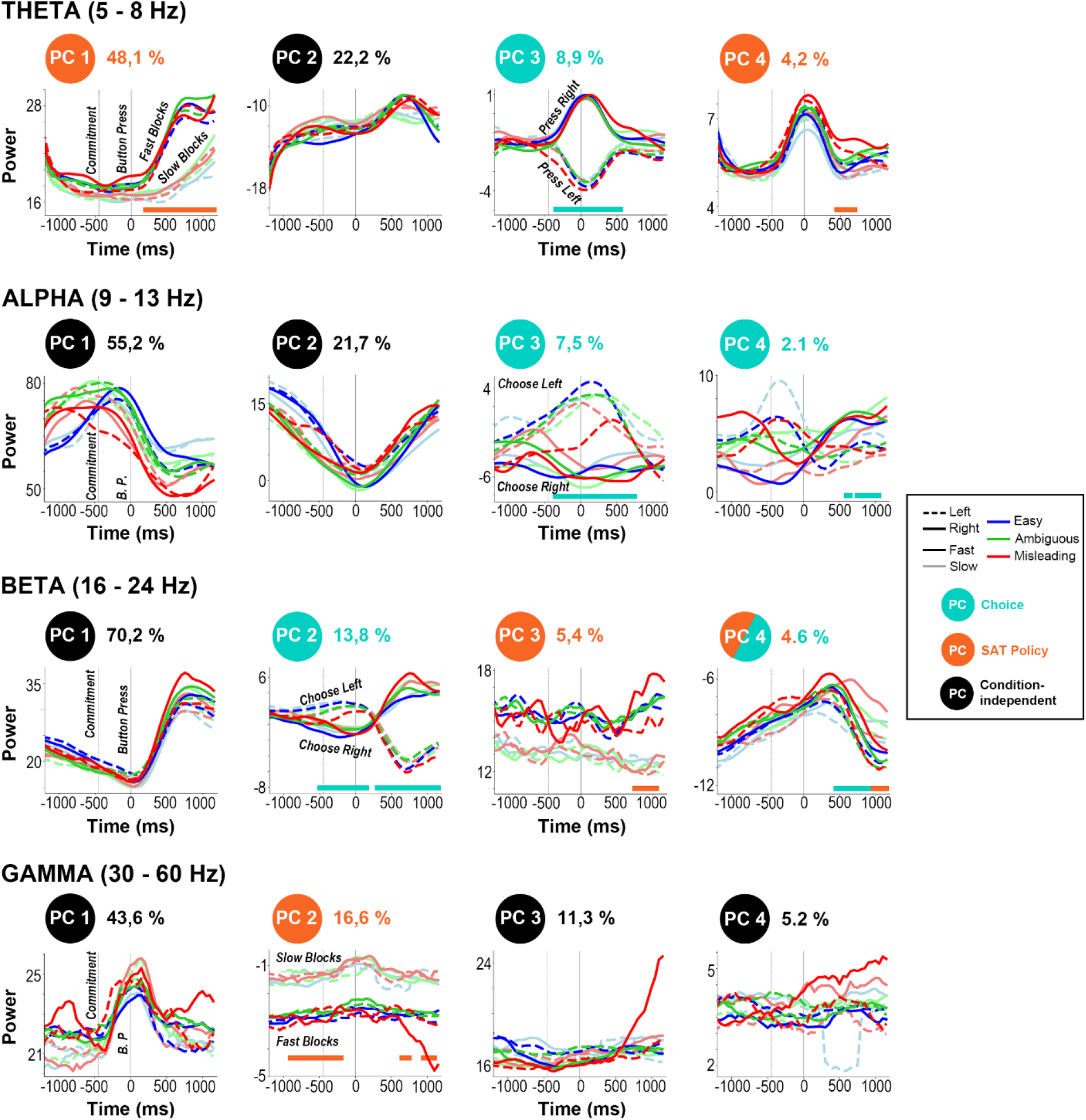
First four principal components (PCs) computed in theta (4 – 8 Hz), alpha (9 – 13 Hz), beta (16 – 24 Hz) and gamma (30 – 60 Hz) frequency bands for easy (blue), ambiguous (green) and misleading (red) trials during fast (bright) and slow (dim) blocks, and for left (dashed) and right (solid) choices on data time-locked to button press (from -1250 to +1200 ms). In each panel, the second vertical dotted line indicates button press, and the first indicates the estimated moment of commitment (i.e., Decision Time, estimated as Button Press Time minus the mean Reaction Time across subjects in the Delayed Response task), 448ms earlier. Percentages next to PCs indicate the amount of variance explained, and PC circular labels are color-coded based on whether they significantly discriminate between task-related conditions using an LDA classifier: choice made (left vs. right, turquoise), SAT policy (fast vs. slow block, orange), and condition-independent (black). Colored horizontal lines mark time intervals with significant effects.

### Alpha and beta patterns of neural activity track sensory evidence during deliberation

First, we sought to identify principal components of neural activity involved in tracking sensory evidence over the course of deliberation. Out of the top 20 PCs computed for each frequency band of interest, we found that time courses (locked on button press) of Alpha PC3 (7.5% of explained variance) and Beta PC2 (13.8% of explained variance) were significantly correlated with the temporal profiles of success probabilities (see Methods) associated with easy (blue), ambiguous (green), and misleading (red) trials for right (solid) and left (dashed) choices (Alpha PC 3: *p < 0*.*05 for 5/6 profiles*, mean R = 0.78, std = 0.3 & Beta PC 2: *p < 0*.*05 for 5/6 profiles*, mean R = 0.83, std = 0.2, see Figure 3B and Supplementary Figure S8). As shown in Figure 3, in easy trials, strong sensory evidence (i.e., high success probability) is reflected by a quick increase in alpha PC3 and beta PC2 activity towards the subjects’ ultimate choice (up for left choices, down for right choices). In ambiguous trials, these PCs fluctuate and then slowly increase towards the chosen target. In misleading trials, they initially ramp up towards the opposite target before they switch, around the same time in both frequency bands, to predict the correct target choice. Results from LDA classification between right and left choices also show that alpha PC3 and beta PC2 profiles could significantly predict the participant’s choice from 350ms and from 500ms before the button press, respectively (left vs right, *p < 0*.*05, corrected across time*). Both PCs were thus identified as “choice components” (turquoise color code, Figure 3C, E).

**Figure 3.**
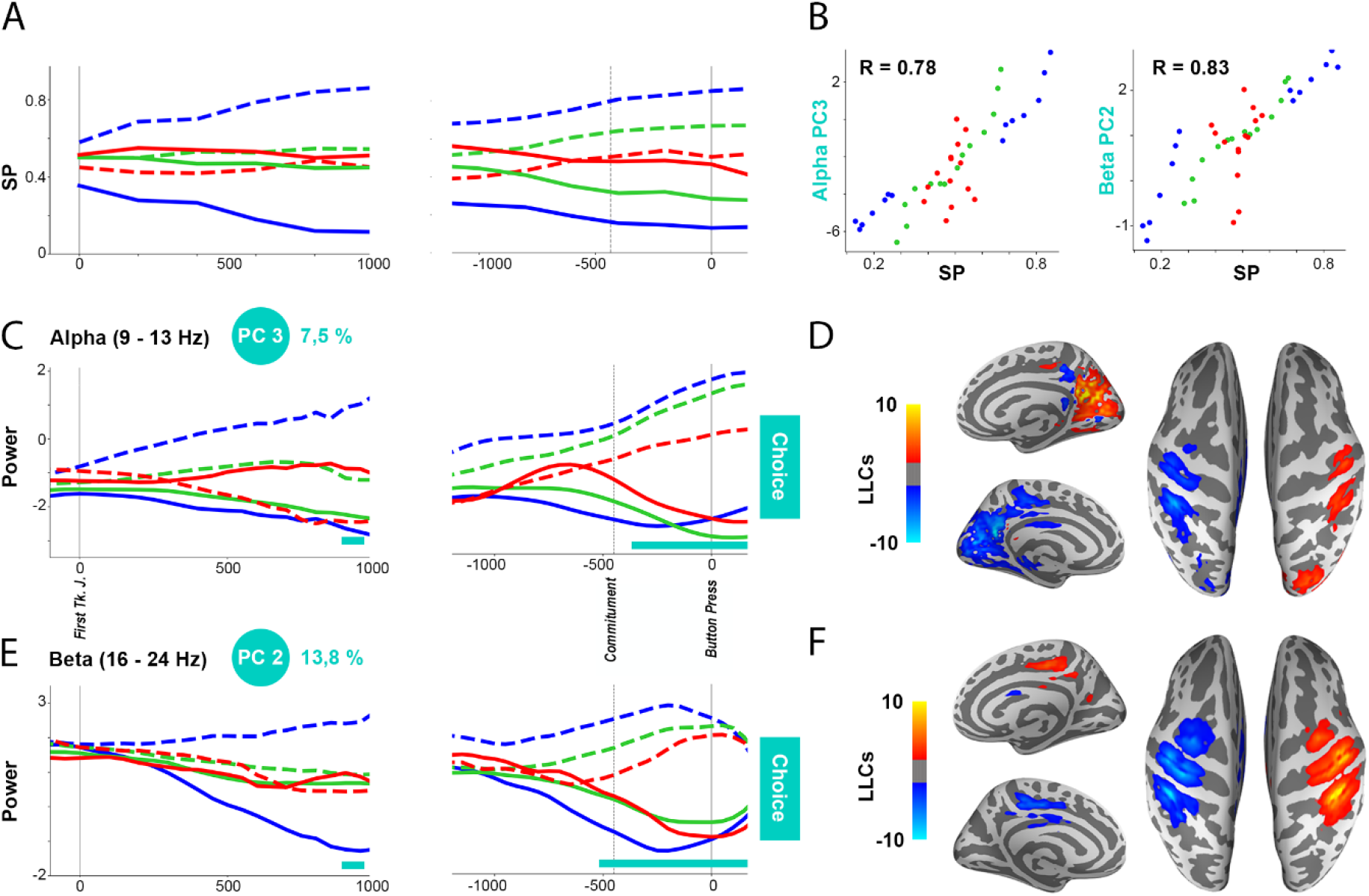
Neural activity tracking sensory evidence during deliberation. **A**. Mean success probability that the target on the left is correct, across subjects and trial types (easy, ambiguous, and misleading trials), for left (dashed lines) and right choices (solid lines) locked on first token jump (first column) and button press (second column), computed every 200 ms. **B**. Scatter plots of mean SP versus mean alpha PC3 (left panel) and beta PC2 (right panel) locked on button press in easy (blue), ambiguous (green), and misleading (red) trials, for left and right choices, for each 200ms time bin. The correlation coefficient is R=0.78 (alpha PC3) and R=0.83 (beta PC2), p-values were below 0.05 for 5/6 profiles (see Supplementary Figure S8 for more details). **C**. Principal component alpha PC3 locked on first token jump and button press during easy, ambiguous, and misleading trials for left (dashed lines) and right choices (solid lines). Horizontal lines mark time intervals with significant effects of choice (turquoise, left vs right), assessed by a Linear Discriminant Analysis (LDA) algorithm (using permutations corrected with maximum statistics at *p < 0*.*05*). **D**. Leading loading coefficients projected on 3D inflated MNI brains for alpha PC3. **E, F**. Same as **C** and **D**, for beta PC2.

To investigate which brain areas contributed most to these PCs, we projected the loading coefficients (i.e., weights computed in the PCA analysis) involved in generating those PCs onto 3D MNI brains in the MEG source space (see Figure 3D, F). We then used the HCPMMP1 atlas^55^ to automatically assign sources to brain regions of interest (ROIs) and highlight similarities and differences between brain regions generating the two PCs tracking the state of sensory information to inform action choices. We show a clear lateralization of the top 5% of loading coefficients across all sources (referred to as “Leading Loading Coefficients, LLCs”, see Methods as well as Supplementary Figures S6 and S7 to see Leading Coefficients and LLCs for the top 4 components of each frequency band, respectively) involved in generating both alpha PC3 (1073/8096 sources) and beta PC2 (860/8096 sources). Moreover, we found that beta PC2 was mostly generated by somatosensory and motor areas including Brodmann area 2, 3b, 4 and the anterior intraparietal area (AIP, see Supplementary Table S2), as well as premotor and superior parietal areas. On the other hand, alpha PC3 was mostly generated by the posterior cingulate cortex, somatosensory and motor areas, but also included primary and early visual cortices (see Supplementary Table S1).

Of note, theta PC 3 was also identified as a choice-related component (see Figure 2) but it showed no significant correlations with success probabilities during the period of deliberation (mean R = -0.01, std = 0.8, *p>0*.*05 for 6/6 profiles*). Instead, theta PC3 peaked around the time of the button press, and the spatial distribution of its LLCs was found in the somatosensory, motor (35 % of LLCs, see Supplementary Table S1) and premotor (16%) cortex, advocating for their strong involvement in motor execution (button presses) once a decision had been made.

### The role of alpha and beta neural trajectories in commitment and button press

To help us further understand the functional role of alpha and beta PCs during dynamic decision-making, we plotted 3D trajectories of the evidence-related components alpha PC3 and beta PC2 against the top two components of each band (Figures 4 and 5), excluding SAT policy components (beta PC3). These were plotted for the different trial types (easy, ambiguous, and misleading) for left and right choices, and averaged across fast and slow blocks (to see trajectories for fast blocks and slow blocks independently, see Supplementary Figure S9). Figure 4A shows two views of the alpha band trajectories (see animated alpha trajectories in Supplementary Video S1), with the individual PCs shown in Figure 4B, and their respective leading loading coefficients (LLCs) in Figure 4C. We found that the first alpha PC was condition-independent and increased until it reached a peak before movement execution (button press), and its LLCs were located in the posterior cingulate cortex and the primary visual area V1 (63 % and 14% of LLCs, see Supplementary Table S1). Alpha PC2 showed a condition independent decrease in activity until button press and was mostly generated by somatomotor (areas 2, 3 and 4, 32% of LLCs) and posterior cingulate cortex (21 % of LLCs, see Supplementary Table S1). Finally, as noted above alpha PC3 tracked sensory evidence and was generated from the posterior cingulate cortex, somatosensory and motor areas, primary and early visual cortices, and the dorsal visual stream. We then showed that during the deliberation period, trajectories projected in a 3D neural state space track sensory evidence (success probabilities of easy, ambiguous and misleading trials, see previous paragraph), and unfold in what we call the “deliberation subspace”^49^ (PC2/PC3), from bottom to top as time elapses and from left to right based on sensory evidence along alpha (Figure 4A, left panel). Importantly, we found that the moment of commitment (colored circles) corresponds to a sharp bend in alpha neural trajectories, after which they rapidly leave the deliberation subspace and move onto the PC1/PC2 subspace in which they evolve in a clockwise manner (arrows in Figure 4A, right panel). Notably, alpha neural trajectories do not bend at the time of button press (colored triangles in left panel) and are not sensitive to movement execution per se, since they only turn back towards the starting point > 500 ms after movement execution was completed (Figure 4A, right panel).

**Figure 4.**
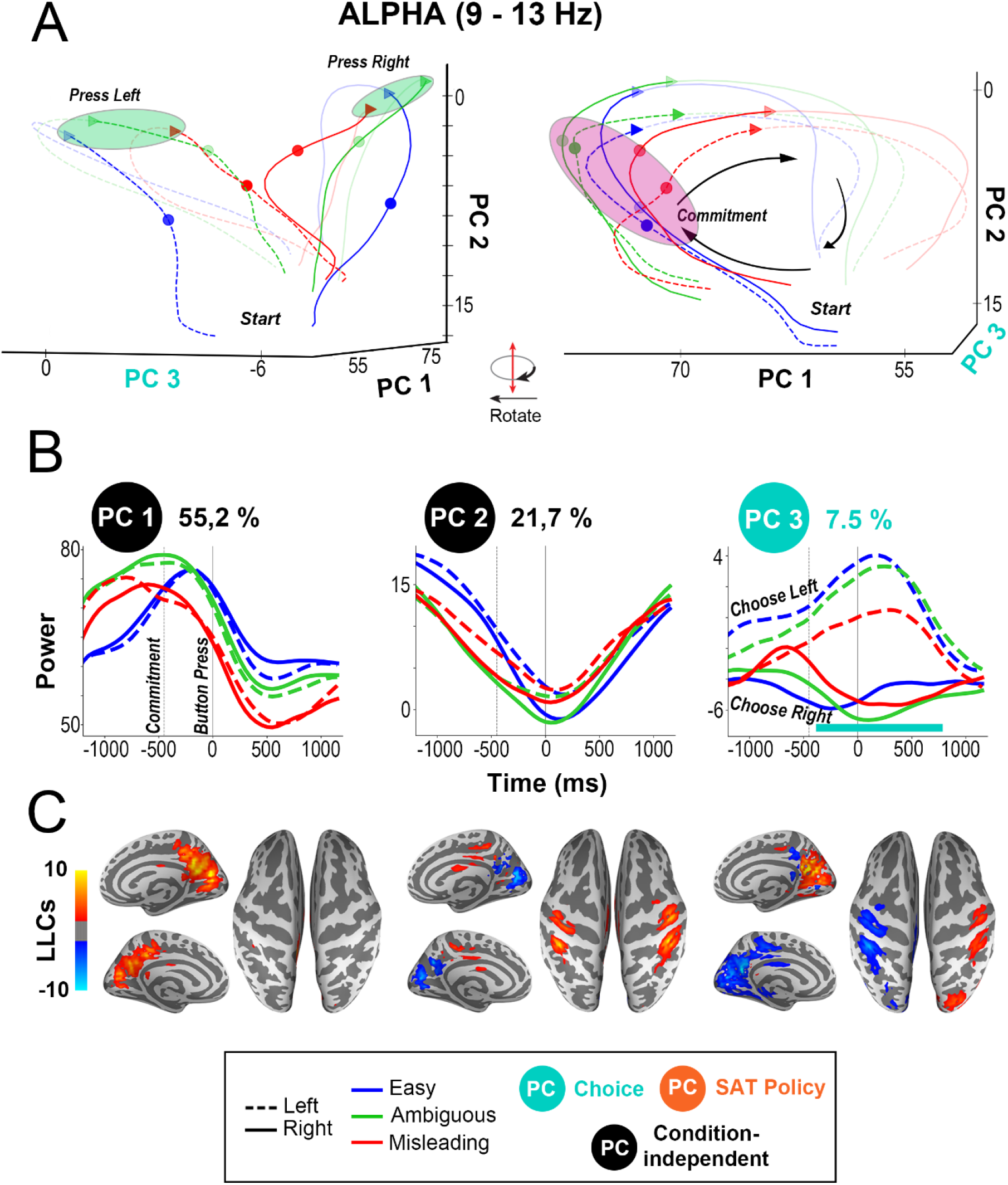
Neural trajectories of alpha PCs in neural state space. **A**. Neural trajectories of alpha PCs 1, 2 and 3 locked on button press [-1200 to 1200 ms], for left or right choices in easy, ambiguous, and misleading trials, averaged across fast and slow blocks. The right panel shows a 90 degrees rotation of the left panel. Small colored circles indicate the times at which commitment occurs, and small colored triangles indicate the time of button press for each individual trial type. Black arrows indicate how neural trajectories unfold over time. Purple ellipses indicate the region in which commitment occurs (indicated for individual trial types by small colored circles) and green ellipses indicate the time of button press (indicated for individual trial types by small colored triangles). **B**. Principal components Alpha PC1, 2 and 3 locked on button press during easy (blue), ambiguous (green) and misleading (red) trials for left (solid lines) and right choices (dashed lines). Horizontal lines mark time intervals with significant effects of choice (turquoise, left vs right) and SAT policy (orange, fast vs slow blocks), assessed by a Linear Discriminant Analysis (LDA) algorithm (using permutations corrected with maximum statistics at p < 0.05). **C**. Leading loading coefficients projected on 3D inflated MNI brains for PC1, 2 and 3 in the alpha band.

**Figure 5.**
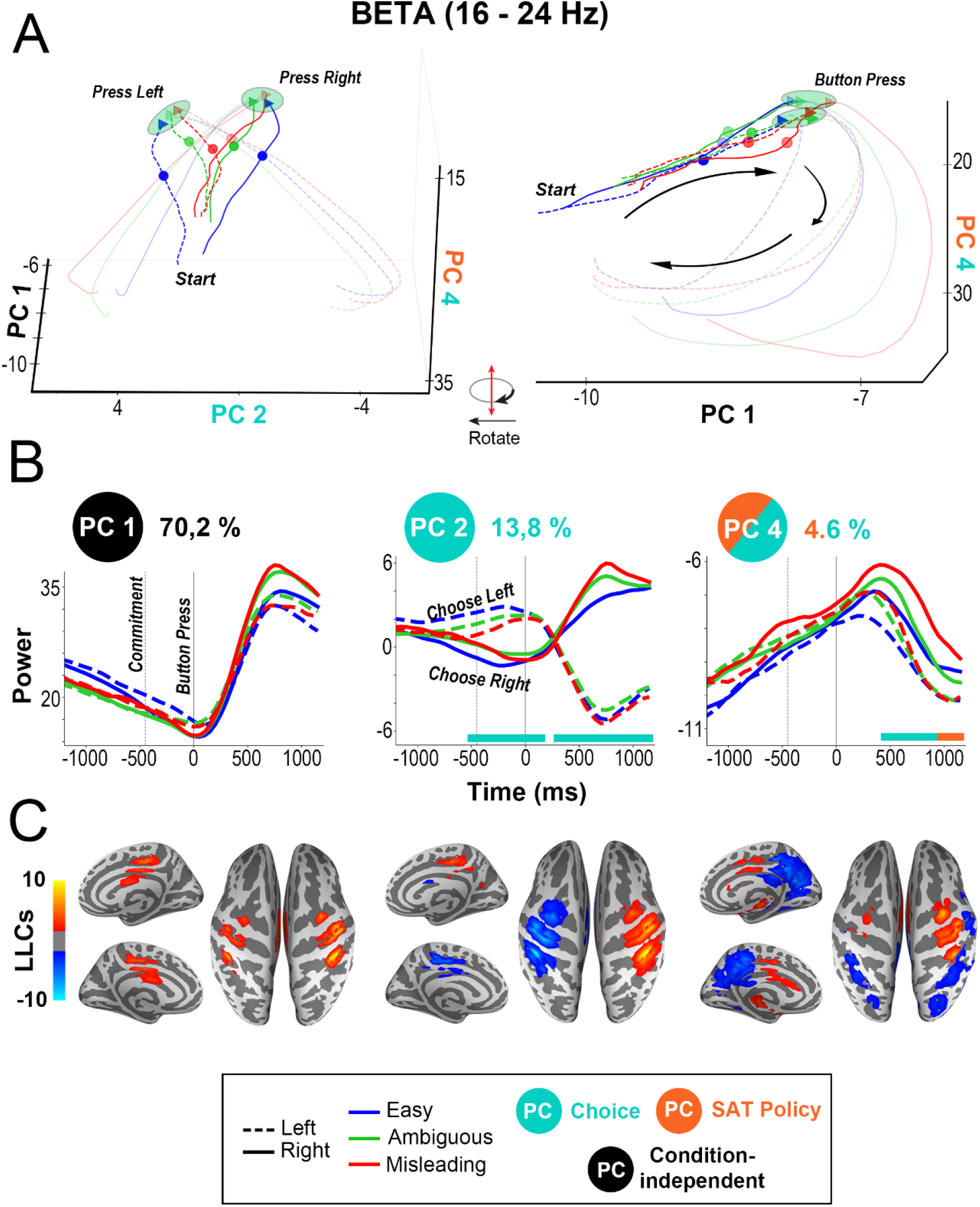
Neural trajectories of beta PCs in neural state space. Same as Figure 4, but for beta PCs 1, 2 and 4.

In contrast with the neural trajectories of alpha PCs, which bend around the time of commitment, the neural trajectories of beta PCs only bend at the time of button press, regardless of the task condition (see animated beta trajectories in Supplementary Video S2),. After button press, beta neural trajectories cross and separate between right and left choices, before turning back in the direction of the starting point (Figure 5A). We found that beta PC1 activity was condition independent and exhibited a gradual decrease before button press, reaching a (negative) maximum around button press onset. After button press, beta PC1 activity transiently increased (i.e., “rebounded”) and peaked around 600 ms (see Figure 5B, left panel). Figure 5C shows that beta PC1 LLCs were mostly located in somatosensory motor regions (areas 2, 3a, 3b and 4, 42%, see Supplementary Table S1 and S2) and in the anterior part of the posterior cingulate cortex (areas 23c and 23d, 13%). Importantly, the beta PC1 temporal profile and LLCs are consistent with the extensive literature describing the typical patterns of beta oscillations before (i.e., gradual decrease), during (negative peak) and after (beta “rebound) voluntary movements^56^. Beta PC2 was found to significantly discriminate both left vs right decisions (i.e., choice component, *p < 0*.*05, corrected across time*), and was mostly generated by somatosensory and motor areas including Brodmann area 2, 3b, 4 and the anterior intraparietal area, as well as premotor and superior parietal areas. Finally, beta PC4 discriminated both left vs right choices and fast vs slow blocks, but only after the button press. Beta PC4 LLCs mostly originated from the posterior cingulate cortex (21% of LLCs), but also included somatosensory motor (11 %), subcortical regions (10%) and the inferior and superior parietal cortex (10% and 8%, see Supplementary Table S1).

### Effect of speed-accuracy trade-off policy on subcortical brain structures

Lastly, we looked for components that reflected adjustments in the subject’s speed-accuracy trade-off policy. To do so, we used an LDA classifier to identify PCs that significantly discriminated fast and slow blocks for each frequency band. This procedure showed that theta- PC1, beta-PC3, and gamma-PC2 were identified as “SAT policy” components (fast vs slow, p < 0.05, corrected across time, see Figure 6). Beta PC3 and gamma PC2 displayed similar temporal profiles and reflected a shift in power distinguishing between the slow and fast blocks (even before the start of the trial, as shown in Supplementary Figure 3). Based on our LLCs analysis, we found that beta PC3 and gamma PC2 mostly originated from subcortical regions (see Figure 6 and Supplementary Table S1). On the other hand, theta PC1 only discriminated fast and slow blocks after button press onset, and their LLCs were located in subcortical brain structures (38.7 % of LLCs, see Supplementary Table S1), but also in the anterior cingulate cortex (15.7%) and in the orbitofrontal cortex (15.4%). To further understand the neural underpinnings of SAT policies in humans, we computed the source reconstructed MEG activity locked on the first token jump in the volume space of the following subcortical regions using the Freesurfer Aseg atlas^57^: caudate nucleus, putamen, globus pallidus, amygdala, cerebellum, and thalamus. We performed a PCA analysis independently for each region and in theta, alpha, beta, gamma, and high gamma frequency bands. Using an LDA classifier, we found that significant “SAT policy” principal components were only found in higher frequencies (beta, gamma and high gamma frequency bands), and originated from the cerebellum, globus pallidus, caudate nucleus and amygdala (see Figure 6D).

**Figure 6.**
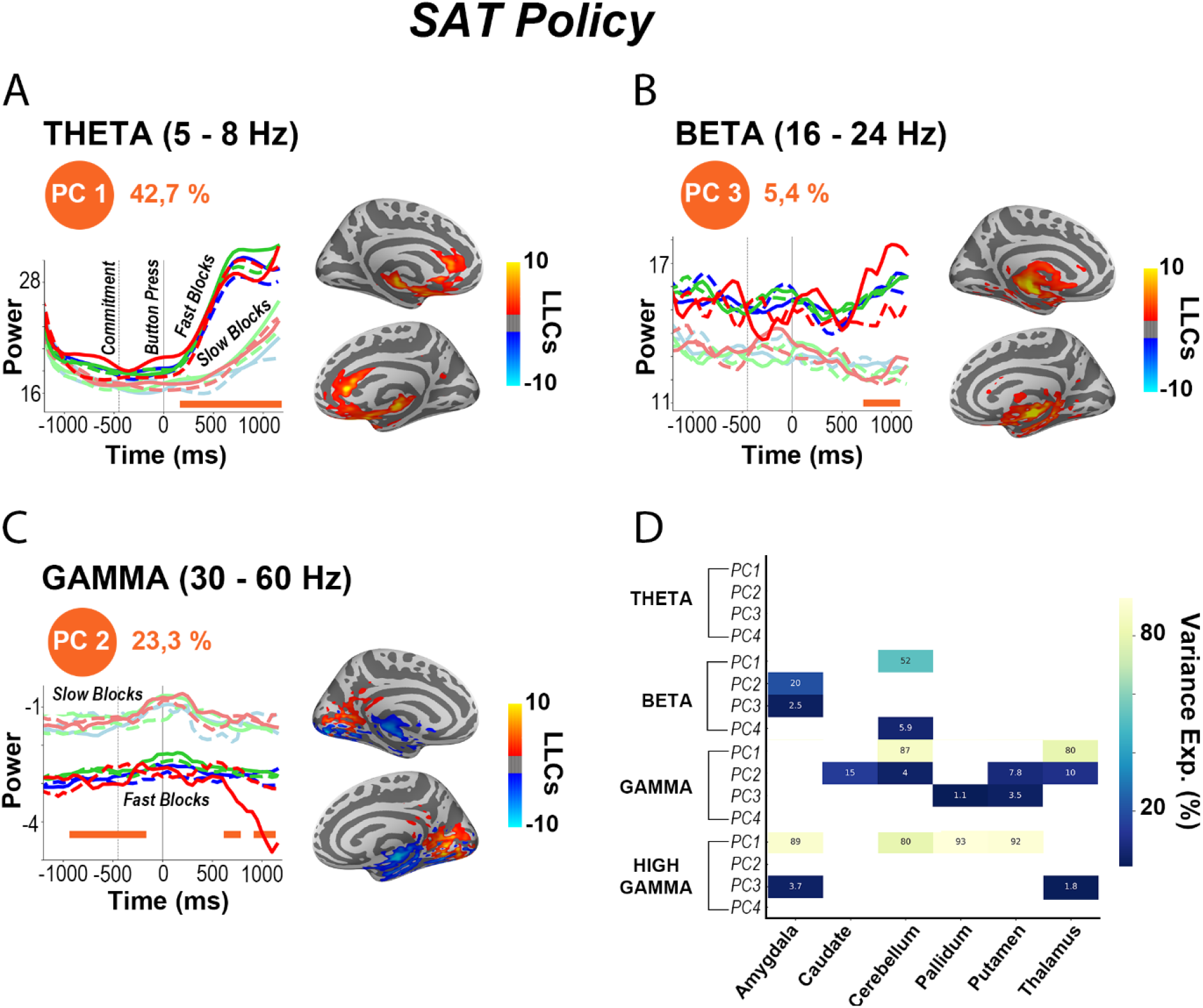
Temporal profiles locked on button press, and their associated Leading Loading Coefficients (LLCs) projected on 3D inflated MNI brains, for the SAT-related principal components: (**A**) Theta PC1, (**B**) Beta PC3 and (**C**) Gamma PC2. Horizontal lines mark time intervals with significant effects of SAT policy (orange, fast vs slow blocks), assessed by a Linear Discriminant Analysis (LDA) algorithm (using permutations corrected with maximum statistics at *p < 0*.*05*). **D**. Heatmap representing significant (fast vs slow blocks, *p < 0*.*05*) SAT policy components for theta, beta, gamma and high gamma frequency bands per region based on the PCA analysis computed on data locked on the first token jump. The PCA analysis was computed independently for each of the following subcortical regions: Amygdala, Caudate, Cerebellum, Pallidum, Putamen and Thalamus for the first 4 PCs. Colors indicate the percentage of variance explained for those components where the SAT effect is significant.

## DISCUSSION

Decisions about actions are arguably the most fundamental kinds of decisions that animals, including humans, face in the natural world. For such decisions, choices are conditioned at least in part by the evidence available in the immediate environment but ongoing changes in the sensory landscape is rather the norm than the exception. However, most decision-making studies have used experimental paradigms in which the informational content of stimuli remains constant over the course of each trial^28^. The present paper provides the first report of neural correlates underlying dynamic decision-making in the human brain using whole brain MEG recordings. Using a dimensionality reduction technique allowed to clearly dissociate and characterize the temporal, spatial and spectral features of neural activity associated with key components of decision-making. In particular, tracking sensory evidence during deliberation was reflected by PCs in alpha and beta bands in the sensorimotor, visual, and posterior cingulate cortex. While alpha PCs activity in visual areas and PCC was associated with commitment to a particular choice, the beta PCs in sensorimotor areas indexed motor execution (i.e., button press). Lastly, we showed the involvement of high frequency PCs (> 16 Hz) generated from subcortical areas (cerebellum, pallidum, caudate nucleus and the thalamus) in speed accuracy tradeoff policies. In this discussion, results will be (1) interpreted in the context of the urgency-gating model and its neural underpinnings in non-human primates^40,46,49,50^ and (2) compared to neurophysiological studies of action decision in humans^1–13^.

### Tracking sensory evidence

First, we showed that principal components (PCs) of activity in the alpha and beta frequency bands were highly correlated with the temporal profile of success probabilities given by the distribution of tokens having jumped in the right and left targets. These “sensory evidence PCs” originated from primary and early visual brain regions, the posterior cingulate cortex, and sensorimotor areas. These results converge with recent human studies using classical decision-making tasks with constant evidence to show that alpha and beta oscillations in visual^9^ and sensorimotor regions^8,58,59^ are crucial for processing sensory evidence. Results are also consistent with accounts suggesting that the PCC could be involved in estimating the need to change behavior in light of new external requirements^60^, including fast perceptual decisions^61^. Importantly, our results also confirm findings about the involvement of parietal, motor, and premotor regions in coding sensory evidence, as reported in non-human primates performing both classical^28^ and dynamic^40^ decision-making tasks. In particular, our results are consistent with studies that used direct single-unit recordings in non-human primates performing the tokens task, which showed the involvement of primary motor (M1) and premotor (PMd) areas in tracking sensory evidence during deliberation and in reaching commitment^40^. However, results did not provide evidence for a role of dlPFC in tracking the state of information given by the tokens’ distribution^49^. As the dlPFC averages across cells with different preferences for right and left responses with no bias for contralateral responses, the lack of results might be due to the fact.

### Alpha oscillations reflect commitment while beta oscillations reflect motor execution

Next, we showed that key decision-making events, namely commitment and motor execution, are reflected by frequency-specific brain activity. We found that the trajectories of alpha PCs in the neural state space “bent” before movement execution and close to the moment of commitment, indicating a role in resolving the competition and committing to a particular choice. This change of trajectories mostly came from alpha PC1, which showed increasing activity until reaching a peak before button press and was mostly generated in posterior regions such as visual cortex and the PCC. These results can be interpreted in the context of literature on the role of alpha oscillations in providing pulsed inhibition, reducing the processing capabilities of given brain areas^62^. In our study, the increased alpha activity in posterior regions could reflect the active and gradual suppression of sensory evidence processing. One possible interpretation is that the peak of alpha activity corresponds to the time of commitment, sensory evidence regions are maximally suppressed to allow a transition of the neural state toward movement preparation and execution.

In contrast, beta PCs did not show a notable change in trajectories at commitment time but instead exhibited a clear change at the button press. These were primarily generated and located in the sensorimotor cortex, suggesting a role in movement preparation and execution. This is consistent with the well-established link between beta frequencies and movement-related processing in the sensorimotor cortex^56^. In particular, the time course of beta PC1 was remarkably similar to the typical decrease in amplitude until movement onset, followed by the increase at cessation of movements, commonly referred to as the “beta rebound”^8^. One hypothesis about the functional role of the beta rebound is that the post-movement period may be used by the sensorimotor cortex to recalibrate or reset the motor system to new conditions, in order to prepare for a subsequent movement^63^. Our findings support this view, as beta PCs in the neural state space clearly show changes of trajectories happening at the time of the beta rebound, redirecting neural activity towards its baseline level, at the start of a trial. Interestingly, our findings support results found in a recent study by Murphy et al. on decision-making in humans using MEG, which highlights both the role of parietal and frontal regions and the importance of visual cortex during action planning^14^. Those authors also reject the idea of perfect evidence accumulation, concluding that recent samples are more strongly weighed than early samples in the decision process as well as in neural activity throughout the visuo-motor system (as in our results). However, in their task the decision timing was externally instructed by a GO cue, and the challenge was to detect changes in the stimulus distribution, whereas in our task subjects were free to commit at any time, giving us a window into their policies for adjusting the SAT and the neural mechanisms of volitional commitment.

### High frequency activity in subcortical regions reflect adjustments of SAT policies

Finally, we found that high frequency (beta, gamma, and high gamma) PCs in subcortical structures reflect adjustment of speed-accuracy trade-off policies. More specifically, we found significant differences between our two block-dependent conditions, encouraging participants toward a more conservative (slow blocks) or risky (fast blocks) SAT policy to optimize their success rate. State-of-the art deep source reconstruction methods suggested that these effects originated in the cerebellum, pallidum, caudate nucleus, amygdala, and thalamus. These observations provide the first experimental evidence that bridges the gap between two streams of research: ***(1)*** studies in which an urgency signal was found in classical random dot motion discrimination tasks when varying the time pressure in humans^10^ and monkeys^64^, and ***(2)*** studies supporting the presence of a neural signature that reflects SAT policies in the basal ganglia of humans^7–10,65^ and monkeys^46^.

### Conclusion

In conclusion, dimensionality reduction of oscillatory power revealed and characterized the neural signatures of key dynamic decision-making processes that allow humans and animals to interact with an environment that can (and does) change continuously over time. Importantly, the use of a dynamic decision-making task made it possible to dissociate activity related to evidence, urgency, and motor preparation, providing new insights consistent with the urgency gating model about the processing of evidence in visual and sensorimotor regions^8,9,28,40,58,59^ (Figure 3), speed-accuracy adjustments involving subcortical areas^7–10,65,66^ (Figure 6), and the process of commitment in motor and premotor cortex^28,32,40,42,47,48^. Our results also cast new light on the functional roles of alpha and beta oscillations in different brain circuits and how they contribute to the commitment to a choice versus its execution (see Figures 4-5). Finally, it is important to note that many of the PCs obtained in our study were remarkably similar to those found when dimensionality reduction was applied to the single-unit data from monkey PMd, M1, dlPFC, and pallidum^49^. This supports the proposal that similar mechanisms are involved when decisions are made by highly trained monkeys versus untrained human subjects, and that the prospects are very promising for unifying the conclusions of single-unit studies in animals with whole-brain human magnetoencephalography.

## ACKOWLEDGEMENTS

KJ is supported by funding from the Canada Research Chairs program and a Discovery Grant (RGPIN-2015-04854) from the Natural Sciences and Engineering Research Council of Canada, a New Investigators Award from the Fonds de Recherche du Québec - Nature et Technologies (2018-NC-206005) and an IVADO- Apogée fundamental research project grant. TT was supported by IVADO Excellence Scholarship – PhD, and a scholarship from FRQS paid by PR in the context of the “Chercheur national – aide à la formation” program. PC received funding from NSERC Grant RGPIN/05245.

## METHODS

### KEY RESOURCES TABLE

| REAGENT or RESOURCE | SOURCE | IDENTIFIER |
| --- | --- | --- |
| **Experimental Models:**<br>**Organisms/Strains** |  |  |
| Humans | N/A | N/A |
| **Software and Algorithms** |  |  |
| Python 3.7 | Anaconda | https://www.anaconda.com/ |
| Matlab 2019b | Mathworks | https://www.mathworks.com/product/matlab.html |
| Labview | NI | http://www.ni.com/fr-fr/shop/labview.html |
| Visbrain | N/A | doi: [10.3389/fninf.2019.00014](https://doi.org/10.3389/fninf.2019.00014) |
| Scikit-Learn | N/A | https://scikit-learn.org/stable/ |
| **Other** |  |  |
| MEG system | CTF | https://www.ctf.com/ |

### LEAD CONTACT AND MATERIALS AVAILABILITY

Further information and requests for resources and reagents should be directed to and will be fulfilled by the Lead Contact, Thomas Thiery.

### EXPERIMENTAL MODEL AND SUBJECT DETAILS

#### Subjects

Thirty-two subjects (aged 18-34, mean age 25.7) participated in this study. 15 were male, and one male was left-handed. Two were excluded before the start of analysis because of large head movement (extent of displacement during one session > 20 mm), and one because of myographic artifacts during MEG scanning. Data from another subject was unusable due to an interruption of the MEG system during the experiment, leaving 28 subjects in total for further analyses. All subjects had normal or corrected-to-normal vision, no history of psychiatric or neurological disorders, and had provided written informed consent prior to the start of the experiment, which was approved by the Research Ethics Committee at University of Montreal, under ethics number CER VN 18-19-05.

#### Behavioral Task

In the MEG scanner, subjects had to perform two behavioral tasks: the “tokens” task and the two choice visuo-motor reaction-time task.

In the tokens task (Figure 1A), 32 healthy subjects between 18 and 34 years old were presented with a central circle and two peripheral target circles (180° apart). At the beginning of each trial, 15 small tokens were randomly arranged in the central circle and began to jump, one-by-one every 200 ms (“predecision” interval), from the center circle to one of the two peripheral targets. The subject’s task was to press a button (left or right) to choose the target he/she believed would ultimately receive most of the tokens. Importantly, the subject could make the decision at any time, and once the choice was reported, the remaining tokens jumped more quickly to their final targets. In separate 5-minute blocks, this “postdecision interval” was set to either 50 ms (in the “fast” blocks) or to 150 ms (in the “slow” blocks). Each subject completed eight blocks, alternating between fast blocks, which encourage hasty behavior, and slow blocks, which encourage more conservative behavior. The task was designed so trials could be classified into three classes of interest: easy, ambiguous and misleading (see next section for more details). Importantly, participants were not given any instruction about speed or accuracy and never told explicitly about block and trial differences, and were only instructed to maximize success rate by making as many correct choices as they could in a given period of time (5 minutes). On each trial, once all tokens had jumped, visual feedback was provided (the chosen target turns green for correct or red for error choices), but there was no feedback or instruction regarding the timing of choices.

In the visuo-motor reaction-time task (two 2-minute blocks before and after the “tokens” task), the 15 tokens originally arranged in the center circle simultaneously jump into one of the two peripheral targets (left or right) after a variable delay. This “GO signal” instructed the subjects to press the button corresponding to the target in which all the tokens have jumped as fast as possible.

#### Behavioral Data Analysis

In the tokens task we can compute, at each moment in time, the success probability *p*_i_*(t)* associated with choosing each target *i*. For instance, if at a particular moment in time the right target contains *N*_R_ tokens, whereas the left contains *N*_L_ tokens, and there are *N*_C_ tokens remaining in the center, then the probability that the target on the right will ultimately be the correct one (i.e., the success probability of guessing correctly) is the following:

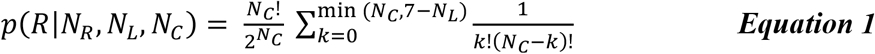

Calculating this quantity for the 15 token jumps allows us to construct the success probability profile *p*_i_*(t)* associated with each trial (Figure 1B).

As far as the subjects knew, the correct target and the individual token movements were completely random. However, to test specific hypotheses about the dynamics of decision making, we interspersed among the fully random trials three specific trial types characterized by particular temporal profiles of success probability. Subjects were not told about the existence of these trials, but 15% of trials were so-called “easy” trials, in which success probability (SP) had to exceed 0.6 after two token jumps, 0.75 after five token jumps and 0.75 after eight token jumps, quickly driving the success probability *p*_i_*(t)* for each toward either 0 or 1. Another 15% of trials were “ambiguous”, in which SP was 0.5 after two jumps, between 0.4 and 0.65 after three token jumps, and then between 0.35 and 0.65 after five jumps, making the *p*_i_*(t)* function stay near 0.5 until late in the trial. Another 10% of trials were called “misleading” trials, in which the SP had to be below 0.4 after three token jumps. The remaining 60% of trials were fully randomized. Thus, the final distribution of trials was as follows: 60% random, 15% easy, 15% ambiguous, and 10% misleading. We used SP to classify random trials into a posteriori classes of trials (e.g., “easy,” “ambiguous,” and “misleading” trials) embedded in the sequence (Figure 1C), yielding an average of 26% easy, 22% ambiguous and 11% misleading trials.

Reaction time in the tokens task was calculated as the time of button press relative to the time of the first token jump. Decision time (DT) was estimated by subtracting from the reaction time the subject’s mean reaction time from the visuo-motor reaction-time task. We could then compute for each trial the duration of a decision as well as its SP, computed using ***Equation 1***, at the time of the decision (Figure 1B).

### QUANTIFICATION AND STATISTICAL ANALYSIS

#### MEG acquisition and preprocessing

Data was acquired from subjects in a seated position using a 275-channel VSM/CTF MEG system with a sampling rate of 2400 Hz (no high-pass filter, 660 Hz anti-aliasing online low-pass filter). Three head positioning coils were attached to fiducial anatomical locations (nasion, left/right pre-auricular points) to track head movements during recordings. Head shape and the locations of head position coils were digitized (Polhemus Fastrak, Polhemus Inc., VT, USA) prior to MEG data collection, for co-registration of MEG channel locations with anatomical T1- weighted MRI. Eye movements and blinks were recorded using 2 bipolar electro-oculographic (EOG) channels. EOG leads were placed above and below one eye (vertical channel); the second channel was placed laterally to the two eyes (horizontal channel). Heart activity was recorded with one channel (ECG), with electrical reference at the opposite clavicle.

We computed 20 independent components (Independent Component Analysis, ICA^67^) from the continuous MEG data filtered between 0.5 and 150 Hz, notch filtered at 60 and 120 Hz, and down sampled to 600 Hz. We identified components capturing artifacts from eye-blinks, saccades and heartbeats based on the correlation of ICA component time-series and ECG/EOG channels. We computed a projector from the identified mixing/unmixing matrices **(Φ/Φ**^**+**^) as **I − ΦΦ**^**+**^ and applied it to the raw unfiltered MEG data to remove contributions from these artifact sources. Subsequently, noisy MEG channels were identified by visually inspecting their power spectrum and removing those that showed excessive power across a broad band of frequencies. The raw data were further visually inspected to detect and exclude time segments with excessive noise e.g., from jaw clenching or eye saccades contamination not captured by any ICA component.

### MEG analyses

#### MEG Source reconstruction

A T1-weighted MRI of the brain (1.5 T, 3D MP-RAGE, 0.8 mm isotropic, sagittal orientation (160), TR= 2200 ms, TE= 2.3 ms; FoV= 256) was obtained from each subject after the MEG session. For subsequent cortically constrained source analyses, the nasion and the left and right pre-auricular points were first marked manually in each subject’s MRI volume. These were used as an initial starting point for registration of the MEG activity to the structural T1 image. An iterative closest point rigid-body registration method implemented in MNE python^68^ improved the anatomical alignment using the additional scalp points. The registration was visually verified. Forward models were generated from the segmented and meshed MRI (decimation = 6, conductivity = [0.3, 0.006, 0.3]) using Freesurfer segmentation pipeline (with default parameter settings^57^) and MNE^68^. The individual high-resolution cortical surfaces (about 120,000 vertices) were downsampled to 8,196 triangle vertices to serve as image supports and define the MEG source space. The epoch data were additionally baseline corrected by subtracting the mean across all time points and epochs for each channel between −200 and 0 ms before the first token jump, and a shrunk co-variance^69^ was estimated across all trials from this time window. The inverse operators were generated using the dynamic Statistical Parametric Mapping method (dSPM, using the Tikhonov regularization coefficient lambda2 = 1), applied at the single-trial level. To perform group analyses, the surface vertices of individual subjects were morphed towards vertices in a reference “fsaverage” space from freesurfer (see surfer.nmr.mgh.harvard.edu/fswiki/fsaverager).

#### MEG Spectral analyses

We first used the FOOOF toolbox^54^, a model-driven method used to parameterize neural power spectra as a combination of an aperiodic component and putative periodic oscillatory peaks of our MEG data. The benefit of using FOOOF for measuring neuronal oscillations is that peaks in the power spectrum are characterized in terms of their specific center frequency, power and bandwidth without requiring predefining specific bands of interest and controlling for the aperiodic component. This method was applied to sensor-level data across all subjects, from - 1500ms to +2000ms around button press, in the frequency range from 1 to 35 Hz (see full report in Supplementary Figure S1). Based on the results given by the FOOOF framework, we chose the following frequency bands: theta (θ) [5–8 Hz], alpha (α) [9–13 Hz], beta (β) [16–24 Hz], and we split high frequency bands into gamma (γ) [30–60] and high gamma (γ) [60–90]. We then extracted the time resolved power by filtering the source reconstructed raw MEG signals using a finite impulse response filtering (FIR1, order = 3) and then computing the Hilbert transform. The signal was then averaged over 500 ms time windows with an overlap of 50 ms. This was computed from 500 ms before to 1000 ms after the first token jump, and from 1500 ms before to 2000 ms after button press, feedback, and decision time. Whenever present, baseline normalization was only used for visualization purposes (time-frequency maps and single trial representation). Baseline normalization was achieved for each frequency band, by subtracting then dividing by the mean of a 400 ms baseline window, i.e., the pre-stimulus rest period ([-500 ms, -100 ms]) before the first token jump.

#### MEG Principal Component Analysis

To calculate the principal components of neural activity, we used the source reconstructed power data computed from 1200 ms before to 600 ms after button press individually for each frequency band. This time window was chosen so it incorporates the token onset movement across all participants and trials. We then grouped trials into twelve non-overlapping classes based on block type (fast or slow), choice (left or right), and trial types (easy, ambiguous or misleading). In each group, power envelopes were averaged together for each frequency band, regardless of whether the choice was correct, or the reaction time was early or late. For each of these twelve classes, we constructed a 28×61×8196 (for the range [-1250; +1200 ms] relative to button press onset) 3D tensor where dimensions correspond to subjects, time bins and sources, respectively. We then concatenated these twelve classes into a 20496 × 8196 matrix (where 20496 = 12 conditions x 28 subjects x 61 time bins and 8196 is the number of source nodes) on which we performed standard Principal Component Analysis (using the pca function in Matlab 2019b), yielding a matrix of weights (“Loading Coefficients”) across all 8196 sources and PCs as well as the variance explained by each PC, for each frequency band. For further analysis, we only kept the top 20 PCs, which explained 95.3% of the total variance across frequency bands (theta: 95.1%, alpha: 97%, beta: 98.1%, gamma: 95.5%, high gamma: 96.5%) though as will be shown below, most of the interpretation will focus on the first four PCs, which explained 84.2% of the total variance across frequency bands (theta: 83.5%, alpha: 86.5%, beta: 94%, gamma: 76.7% and high gamma: 80.3%).

The time courses of PCs for each of the twelve experimental conditions were computed by taking the product of loading coefficients and original power data (locked on the first token jump, on decision time or on button press), for each subject and for each frequency band. To localize the distribution of weights that contributed most to each PC, we projected the top 5% of loading coefficients (“Leading Loadings Coefficients”, LLCs) onto sources of 3D inflated standard MNI brains. Specifically, we created a distribution of the absolute values of all loading coefficients, and only kept top 5% of sources based on their loading coefficient (absolute) value. Importantly, we ran the same procedure using 3% and 10%, and obtained similar results.

#### Correlation analyses

We computed Pearson correlation coefficients (R) and their associated *p-values* between the time-courses of mean success probabilities (SPs) and the time courses of the top 20 PCs for easy, ambiguous, and misleading trials and for left and right choices averaged across fast and slow blocks (6 conditions), for each frequency band separately. SPs and PCs were computed every 200 ms from 0 to 1000 ms after first token jump and from -1200 ms to 0 ms before button press onset (see Supplementary Figure S8).

### Decoding analyses

#### Signal classification

We set out to evaluate whether time courses of principal components derived from the PCA analysis could significantly classify effects of decision (Left vs Right), sensory evidence (Easy vs Ambiguous vs Misleading), and SAT policy (Fast vs Slow) during dynamic perceptual decision making. To this end, we implemented a machine learning framework for classification using matrices where dimensions correspond to subjects (n=28), conditions (n=2 for Left vs Right and Fast vs Slow, or n=3 for Easy vs Ambiguous vs Misleading), and time bins. LDA classifications were repeated for each frequency band (θ, α, β and γ), and for data locked on the first token jump, on decision time or on button press).

Several classification techniques were initially tested for the single feature classification procedure, including linear-discriminant analysis (LDA), k-nearest-neighbor (KNN) and support vector machine (SVM). The classification accuracy results were similar across the three methods. The LDA algorithm (Fisher, 1936) was the fastest and was therefore chosen for this study given the computationally demanding permutation tests used to evaluate classifier performance. In brief, for a binary classification problem, the LDA algorithm tries to find a hyperplane that maximizes the mean distance between the mean of the two classes while minimizing inter-class variance.

#### Decoding accuracy and statistical evaluation of decoding performance

At the group level, the performance of the proposed classification method was evaluated using a Leave-One-Subject-Out (LOSO) cross-validation procedure. This procedure is a special case of k-fold cross-validation, where all individuals except one are used for training, and the classifier is tested on the data from the omitted participant (i.e., test data). This procedure is repeated iteratively, each time leaving a different individual out of the training. The LOSO cross-validation method efficiently uses data and provides an asymptotically unbiased estimate of the averaged classification error probability over all possible training sets^70^. The statistical significance of the obtained decoding accuracies was evaluated by computing statistical thresholds using permutation tests (*p < 0*.*05*). In other words, a null distribution is generated by repeatedly (*n = 1000*) computing the classification accuracy obtained after randomly permuting class labels^71^.

## SUPPLEMENTARY MATERIAL

**https://www.dropbox.com/scl/fi/indsoe08aguvkveuh1csz/00_Table_S1.xlsx?dl=0&rlkey=6ov9zzv22ttzxqm2517oeyzmo**

**Table S1. Leading Loading Coefficients assigned to Regions of interest (ROIs)** using the HCPMMP1 combined atlas^55^.

**https://www.dropbox.com/s/fno2fspnw2cx8f3/00_Table_S2.xlsx?dl=0**

**Table S2. Leading Loading Coefficients assigned to Regions of interest (ROIs)** using the HCPMMP1 atlas^55^.

**Figure S1.**
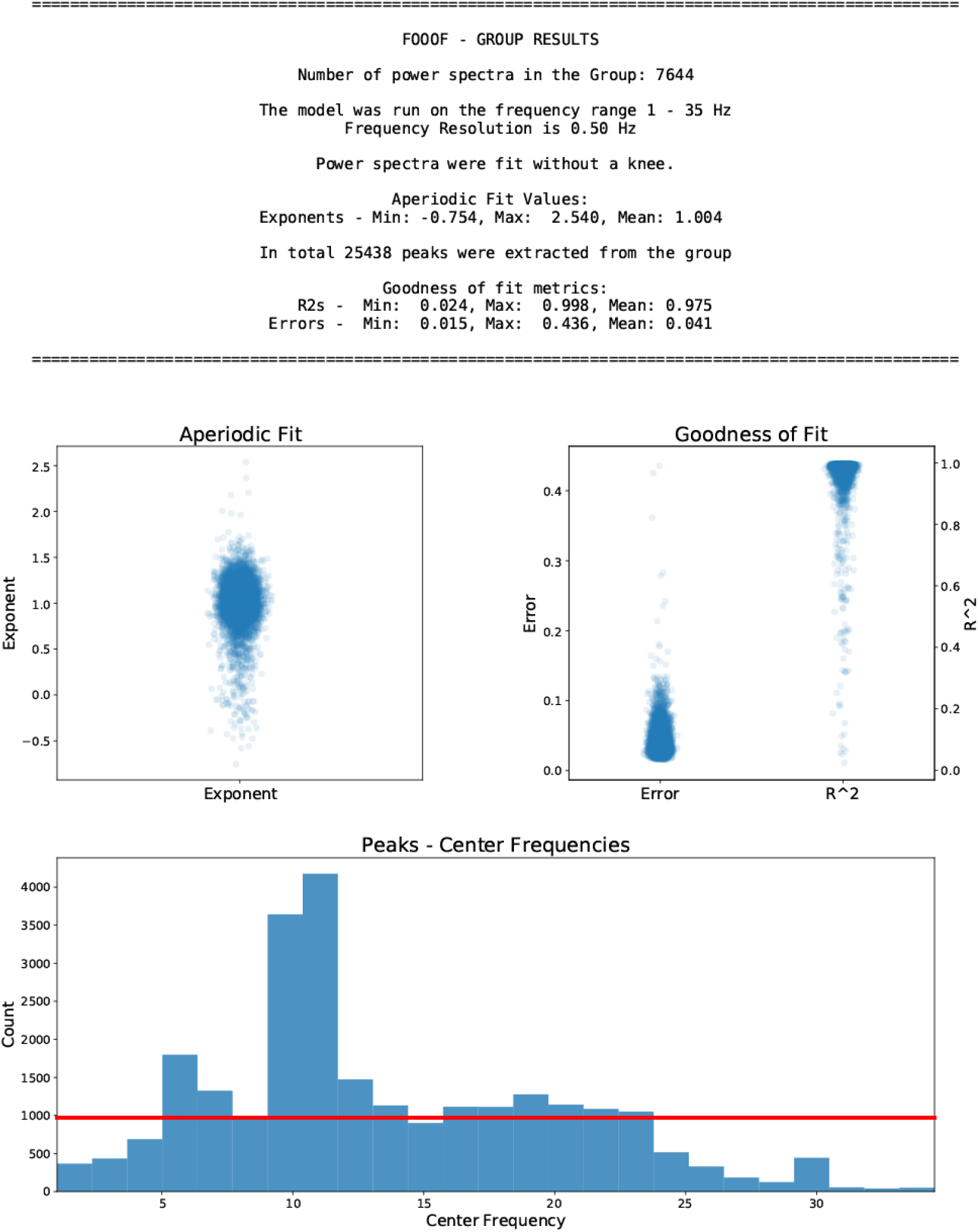
FOOOF toolbox report. To determine the width of the frequency bands for theta, alpha and beta we fit a model on power spectra (n=7644) from all subjects based on the sensor-level data (28 subjects * 273 channels) locked on button press (-1500ms to +2000ms). We chose to group frequencies into three bands: theta [5-8 Hz], alpha [9-13 Hz] and beta [16-24 Hz].

**Figure S2.**
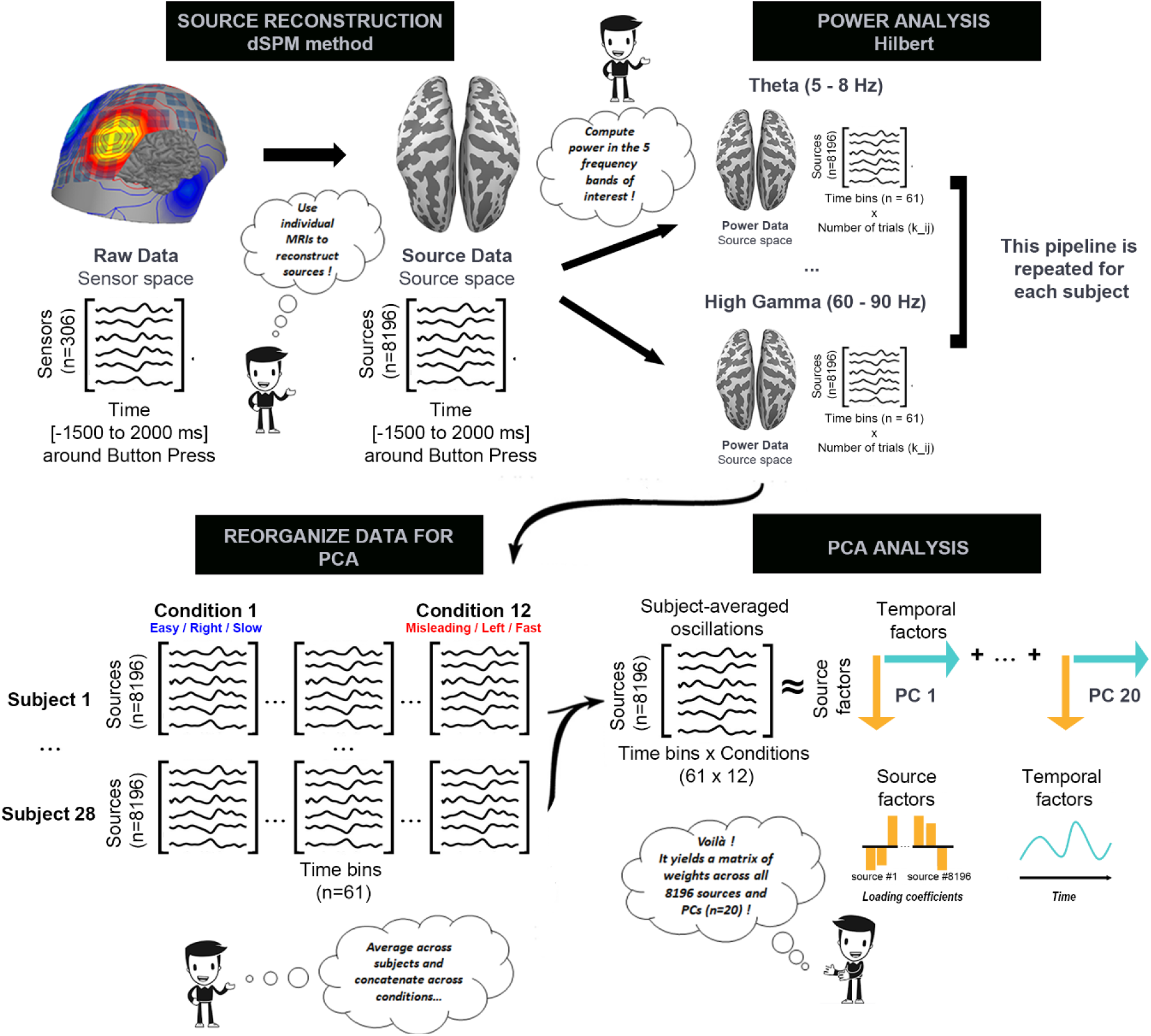
Illustration of the PCA analysis. performed on the source-reconstructed MEG data. We first used the preprocessed raw MEG data in the sensor space combined with the individual T1 MRI images to reconstruct sources. We then computed power envelopes in 5 frequency bands of interest, using the Hilbert method with sliding windows (width, overlap) in single trials, i.e. without averaging envelopes across trials (k represents the number of trials, *i* the condition from 1 to 12, and *j* the subject number from 1 to 28). We then reorganized the data to obtain the following 3D tensor for each frequency band: 12 conditions (averaged across trials in each condition) x 8196 sources (averaged across subjects) x 61 time bins. We then turned this into a 732 × 8196 matrix by concatenating the conditions and performed the PCA analysis to obtain a matrix of weights (“Loading Coefficients”) across all 8196 sources and principal components (PC) as well as the variance explained by each PC, for each frequency band.

**Figure S3.**
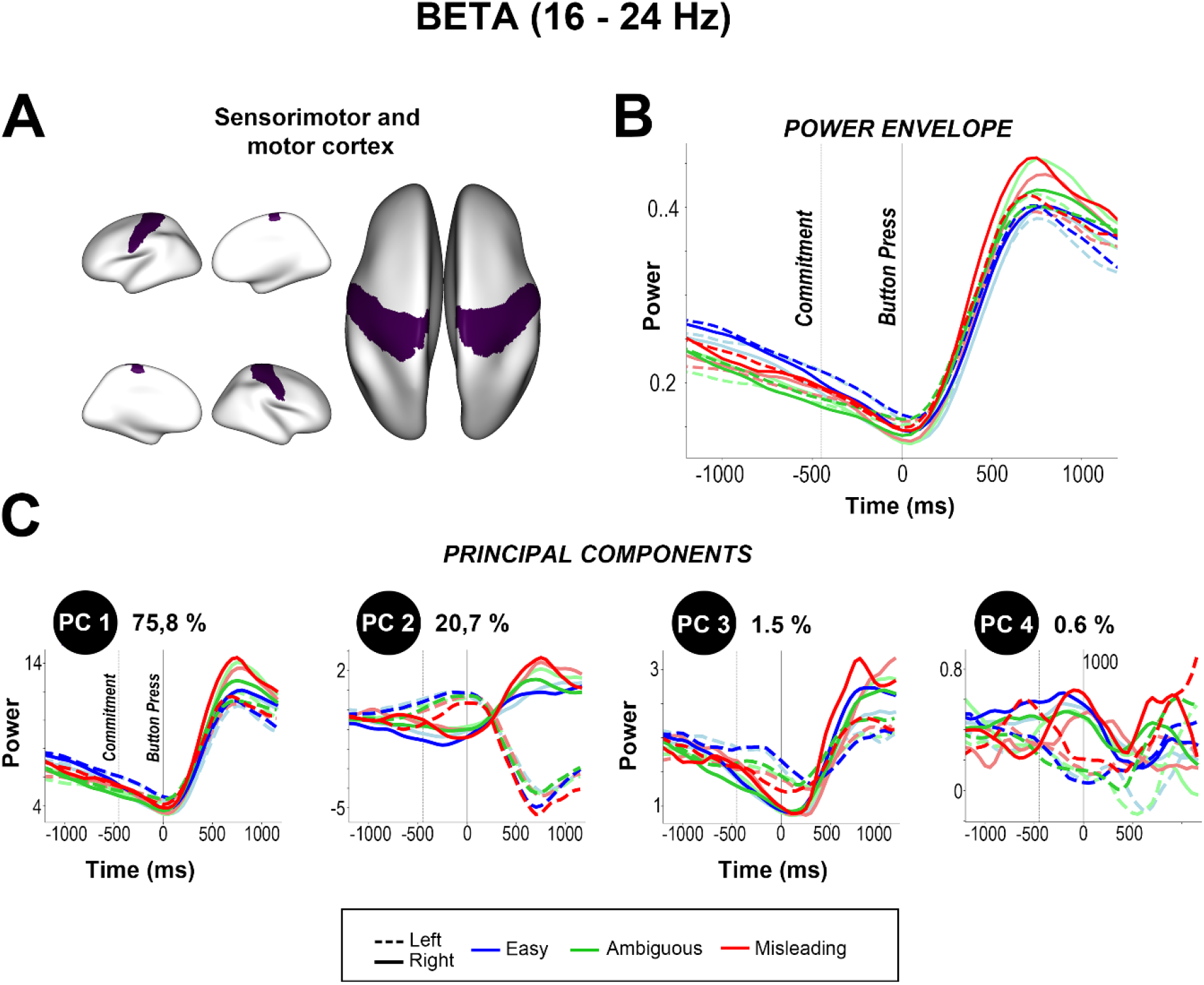
Comparison between beta (16-24 Hz) power envelopes (B) and principal components (C) in the sensorimotor and motor cortex (A), averaged across trials and participants for easy, ambiguous and misleading trials, and for left (dashed) and right choices (solid).

**Figure S4.**
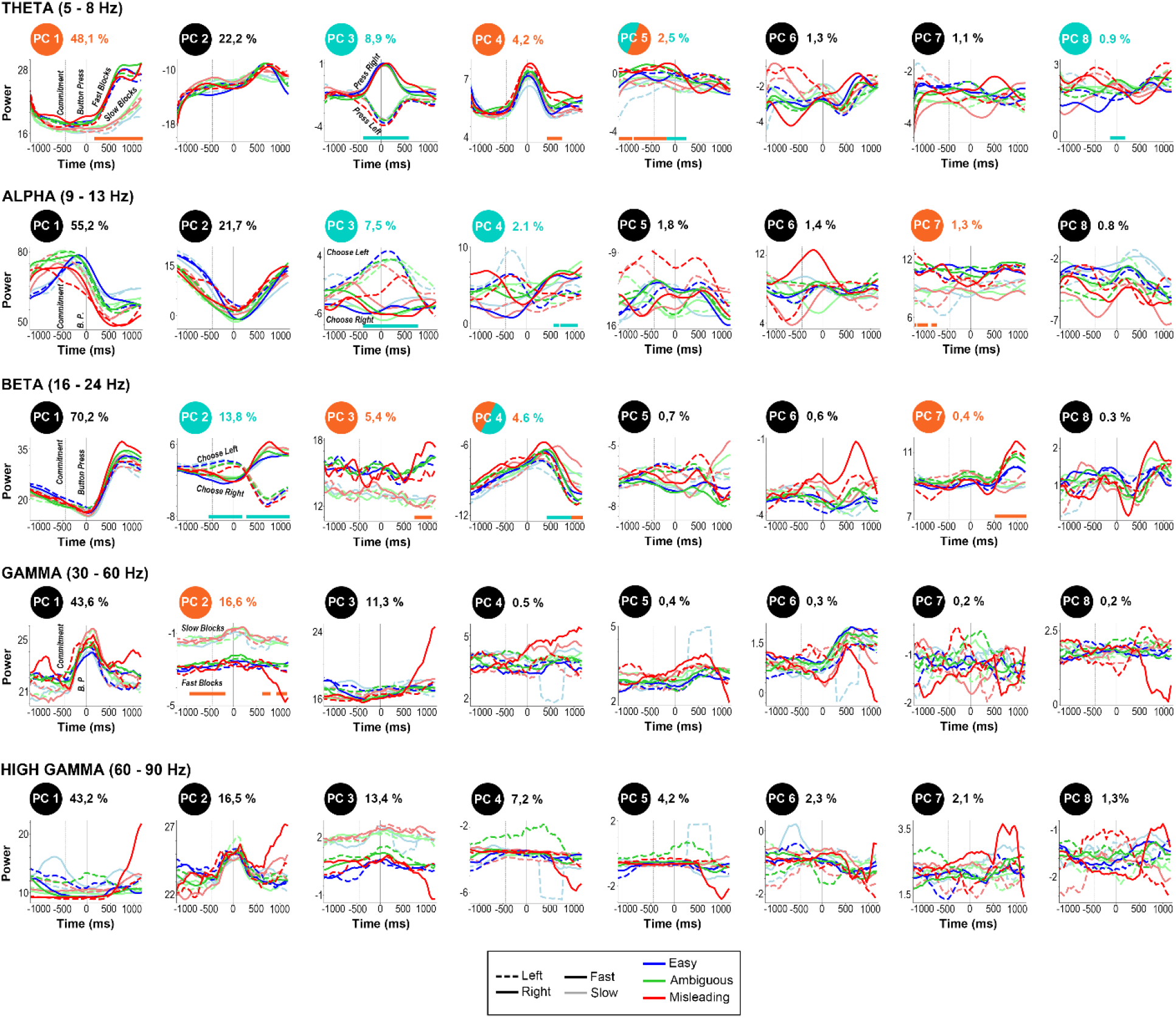
First eight principal components computed in theta (5 – 8 Hz), alpha (9 - 13 Hz), beta (16 – 24 Hz), gamma (30 – 60 Hz) and high gamma (60 - 90 Hz) frequency bands for easy (blue), ambiguous (green) and misleading (red) trials during fast (bright) and slow (transparent) blocks, and for left (solid) and right (dashed) choices on data time locked on button press (from -1250 to 1200 ms). In each panel, the second vertical dotted line indicates button press, and the first indicates the estimated moment of commitment (i.e., mean decision time across subjects obtained in the Delayed Response task), 448ms earlier. Percentages next to PCs indicate the amount of variance explained.

**Figure S5.**
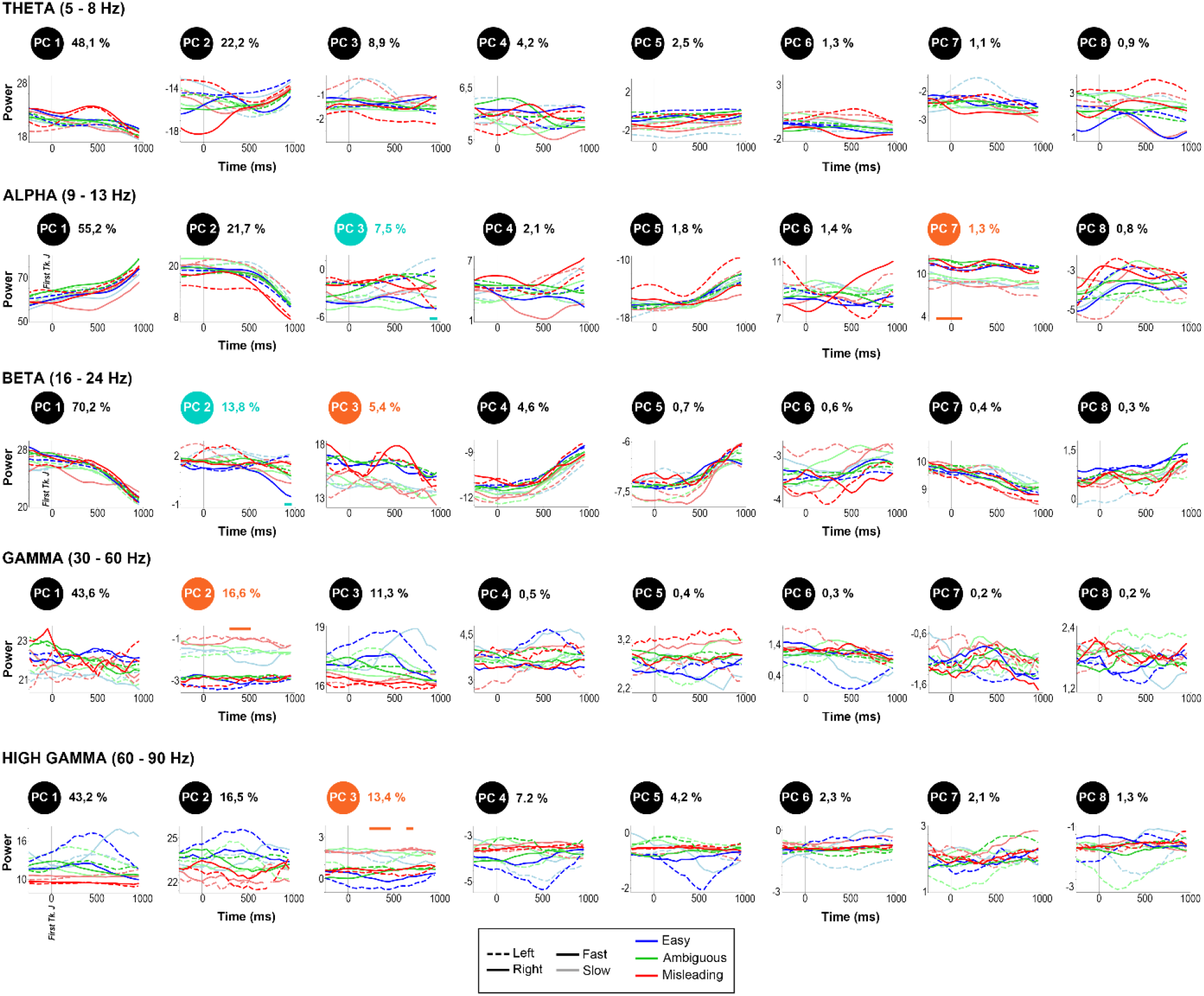
First eight PC components computed in theta (5 – 8 Hz), alpha (9 - 13 Hz), beta (16 – 24 Hz), gamma (30 - 60 Hz) and high gamma (60 – 90 Hz) frequency bands for easy (blue), ambiguous (green) and misleading (red) trials during fast (bright) and slow (transparent) blocks, and for left (continuous) and right (dashed) choices on data time locked on the first token jump (from -200 to 1000 ms). Percentages next to PCs indicate the amount of variance explained.

**Figure S6.**
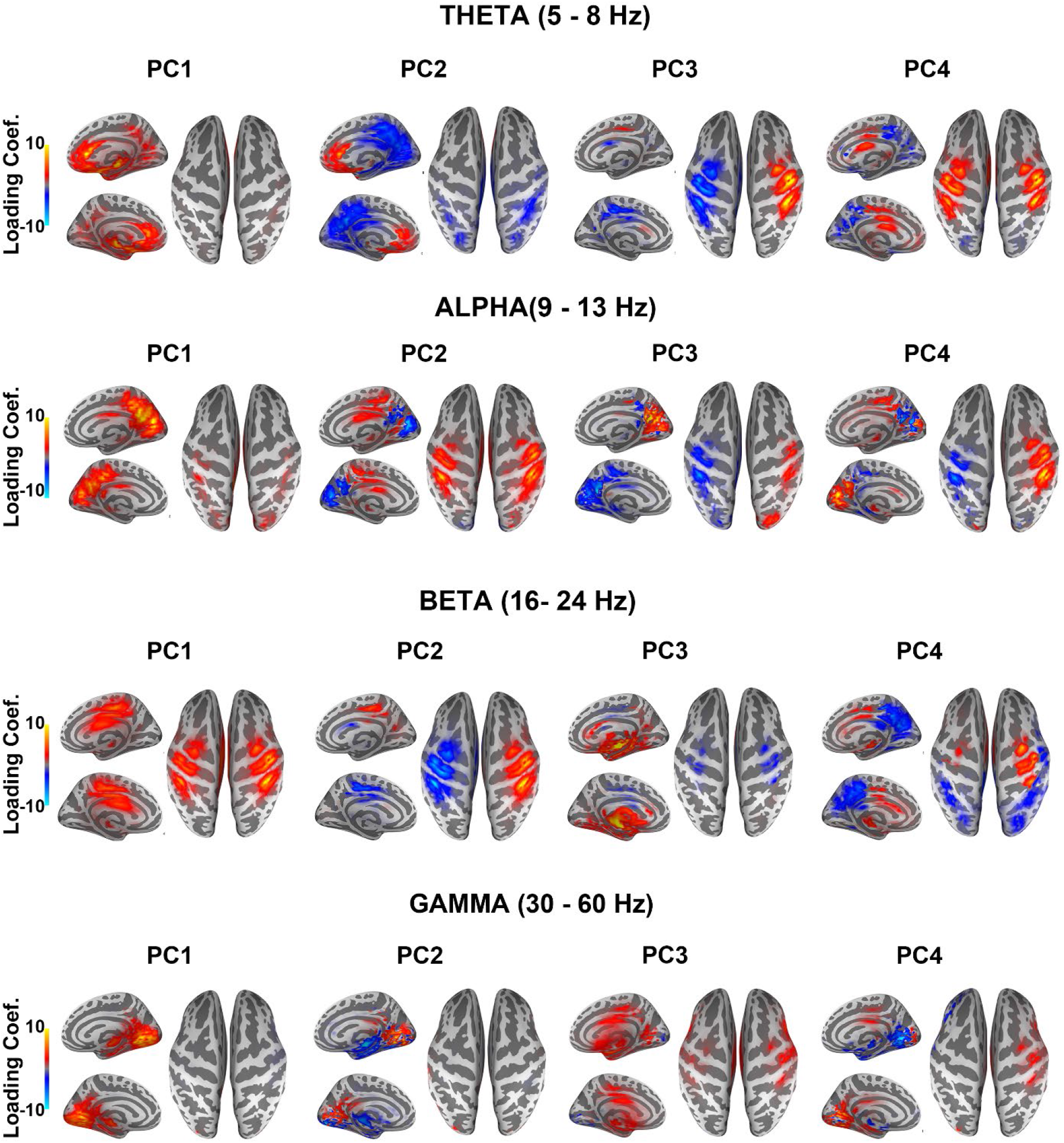
Loading coefficients. projected on 3D inflated MNI brains in the MEG source space for the first 4 components in theta, alpha, beta and gamma frequency bands.

**Figure S7.**
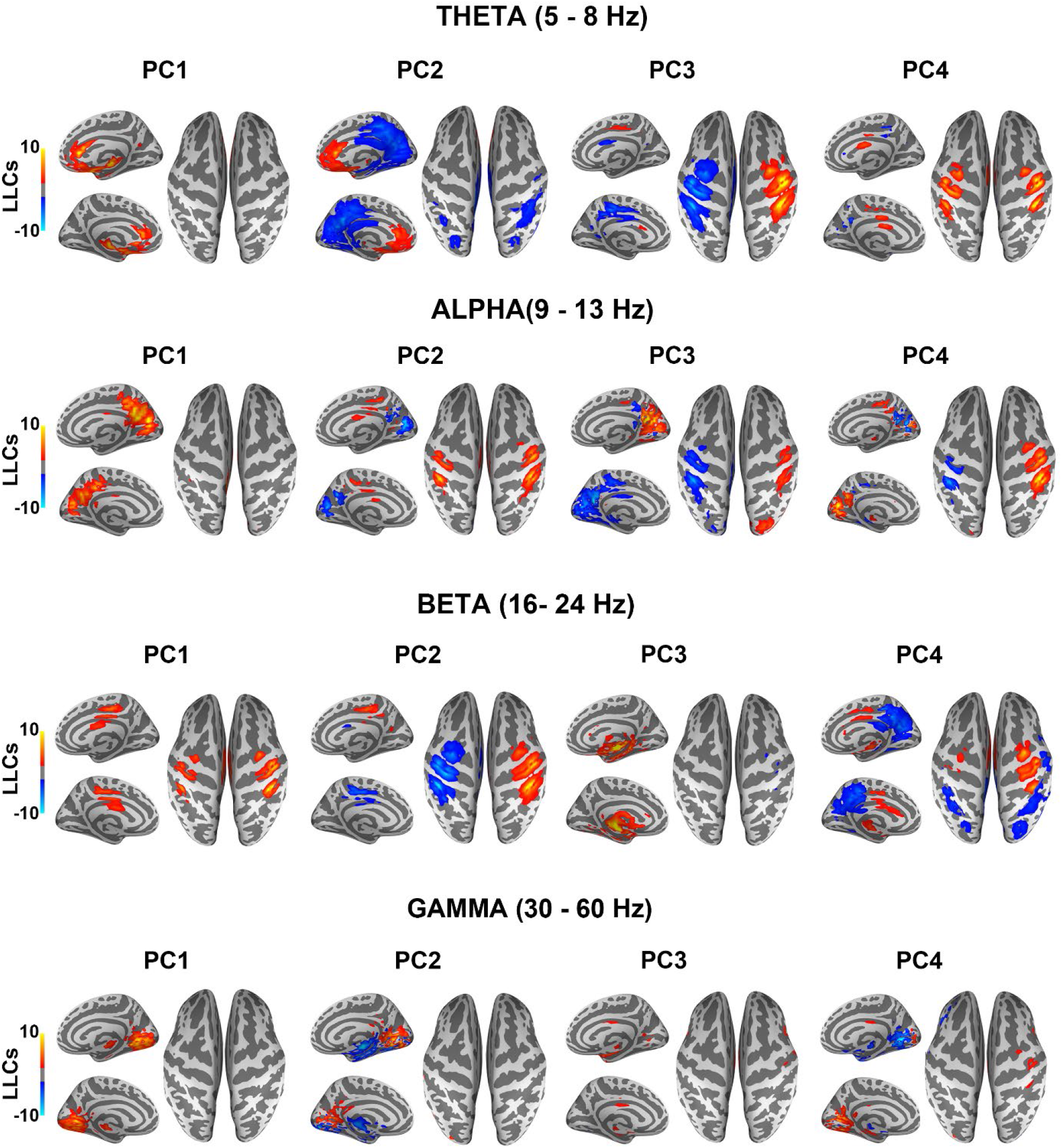
Leading Loading coefficients (LLCs, top 5%) projected on 3D inflated MNI brains in the MEG source space for the first 4 components in theta, alpha, beta and gamma frequency bands.

**Figure S8.**
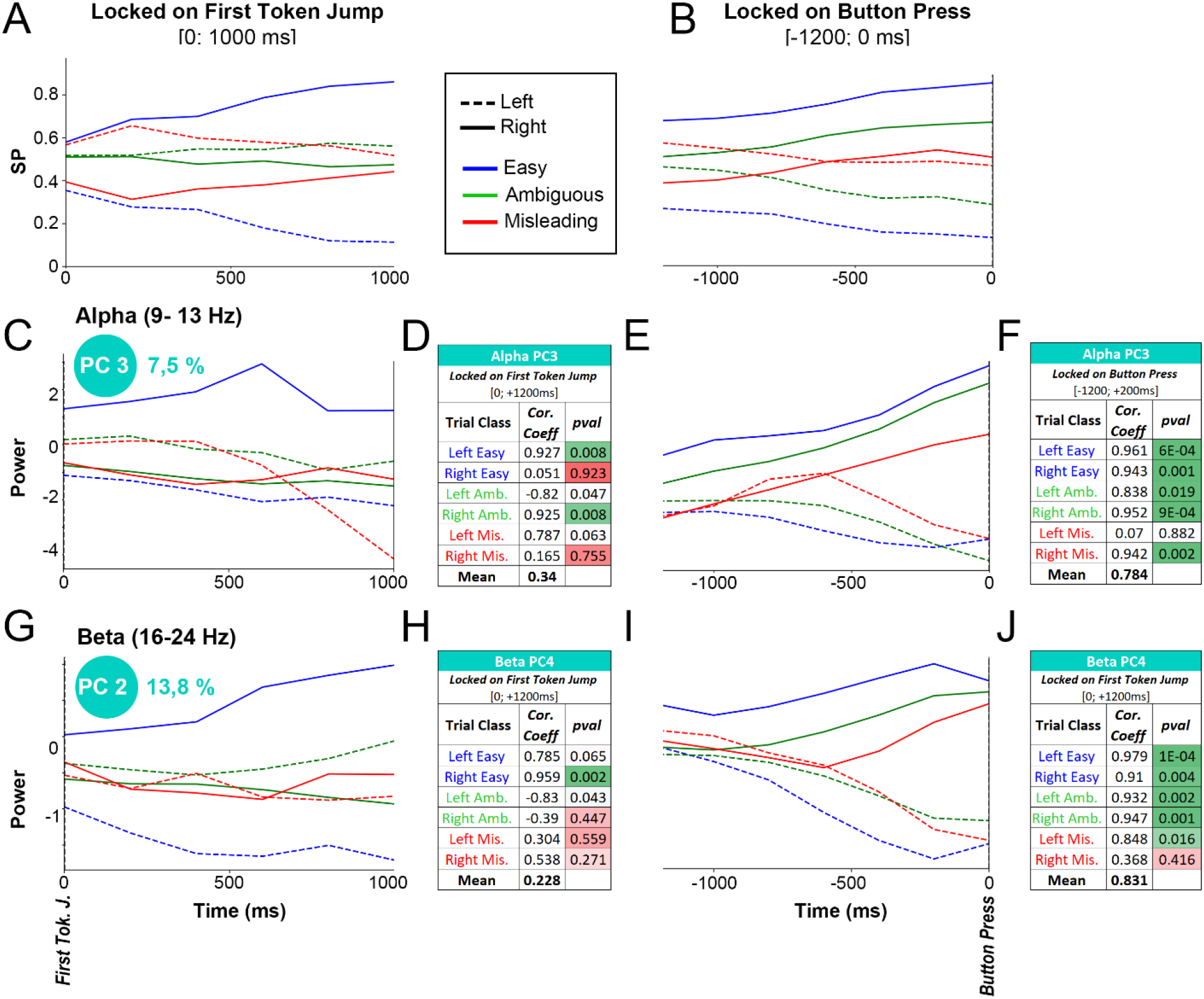
Correlation between success probabilities and PCs. Success probabilities for the target on the right, locked on first token jump [0; 1000 ms] (**A**) and locked on button press [-1200; 0 ms] (**B**). Alpha PC3 and beta PC4 temporal profiles locked on first token jump (**C, G**) and on button press (**E, I**) averaged across fast and slow blocks for easy, ambiguous, and misleading trials and for left and right choices. Tables **D, F, H** and **J** show correlation coefficients and *p-values* between SPs and alpha PC3/beta PC4 locked on first token jump (**D/H**) and button press (**F/J**).

**Figure S9.**
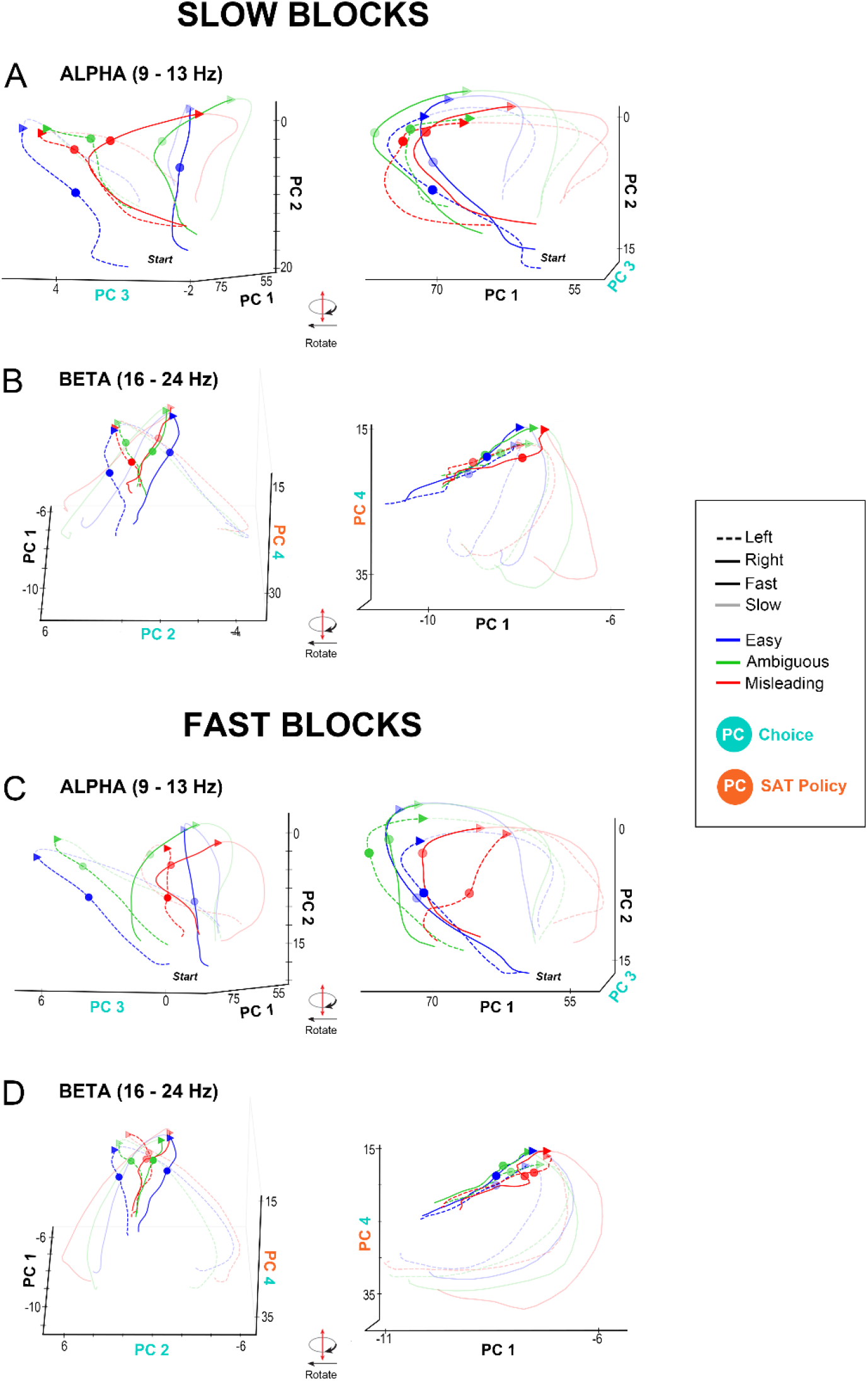
Neural trajectories of alpha and beta PCs in neural state space for fast and slow blocks. Neural trajectories of alpha PCs 1, 2 and 3 and beta PCs 1, 3 and 4, locked on button press [-1200 to 1200 ms], for left or right choices in easy, ambiguous, and misleading trials, for slow (**A, B**) and fast blocks (**C, D**). Small colored circles indicate the region in which commitment occurs, and small colored triangles indicate the time of button press for each individual trial type. Black arrows indicate how neural trajectories unfold over time. The right panel corresponds to a 90 degrees rotation of the left panel.

